# Viral transport in evaporating sessile model respiratory droplets

**DOI:** 10.1101/2025.09.30.679206

**Authors:** Javier Martínez-Puig, Carlos de la Torre Luque, Carolina Arbesu Nieto, Pepijn Hoekstra, Fernando Usera, Beatriz Martín-Jouve, Aránzazu de la Encina, Teresa Bartolomé, Marta Sanz, Fernando Almazán, Ana Oña, Alvaro Marin, Javier Rodríguez-Rodríguez

## Abstract

Viral particles, or virions, may remain infectious in the dry residue of respiratory droplets for times as long as hours. This is counterintuitive, since salt concentration increases dramatically as the drop’s water evaporates, making the drop a harsh environment for virions. It has been hypothesized that the drop components (mainly salt and the glycoprotein mucin) segregate during the evaporation process, with virions being transported away from salt deposits. This would protect them from the salt’s damaging effects. Thus, understanding where virions reside in a drop residue is essential to disentangle the physico-chemical mechanisms that drive their inactivation. However, determining the virion location in an environment as heterogeneous and complex as a respiratory drop is challenging. Here we show experimentally that virions in the drop’s dry residue are found mainly forming aggregates in protein-rich regions, outside salt crystals. The mechanism yielding the observed viral spatial distribution can be found in the internal flow phenomena within the droplet, which are described using numerical simulations. We anticipate our results to be relevant to explain the discrepancies in the infectivity decay rates measured in respiratory drops between previous reported studies.

## 1 Introduction

Many viruses and other pathogens are able to remain infective in the dry residue of a respiratory drop for hours or even days [1]. A priori, this is rather counterintuitive since, as the drop loses its water content to evaporation, the salt concentration inside the drop increases to levels high enough to inactivate any virion present in the drop in a much shorter time scale [1, 2, 3]. Two explanations for this apparent paradox have been proposed in the literature. First, that the presence of organic compounds in respiratory fluids, most notably the glycoprotein mucin, protects virions from the damaging effect of salt ions [4, 5]. Second, that once the water content of the drop is lost by evaporation, salt precipitates into crystals, which self-segregate spatially from virions [1]. This second hypothesis is consistent with the observation that the decay rate of Influenza is higher during the evaporation process than after the residue has dried [3, 6]. However, these findings are contradicted by other authors. For instance, Rockey and coworkers [7] observed that, in some circumstances, an elevated salt (i.e., ion) concentration was linked to a slower inactivation rate for Influenza virus. Moreover, Oswin and coauthors [8] report an abrupt decay in viral viability right after efflorescence (salt precipitation) in micrometric aerosolized droplets.

These examples illustrate the discrepancies found across different studies in the literature [9]. We can conclude that, despite the significant progress on the matter, the precise physico-chemical mechanisms that drive viral inactivation inside respiratory drops are still unknown [10]. A critical missing ingredient to unravel these mechanisms is to know where virions are located inside the dry residue of the droplets as water evaporates [5, 11, 12], and equally important, what is the physico-chemical microenvironment that surrounds them there. Direct observation of the virions inside the drop residue would partly settle this question, but this is not a trivial task. Although there exist many techniques to visualize virions in purified suspensions [13, 14] or in homogeneous, diluted gel-like structures [4, 15, 14], visualizing the distribution of a large number of viral particles in complex heterogeneous structures, like the residue of a respiratory drop [16], is still challenging.

One of the first attempts to visualize the viral distribution within a drop was made by Vejerano & Marr [11]. They used a non-specific fluorescent tagging on the *ϕ*6 phage to localize these virions in the residue of a drop of artificial saliva. They found isolated fluorescent spots compatible with virion aggregates distributed without any specific spatial pattern in the dry residue of the drop. Unfortunately, the limited number of experiments reported and the fact that the fluorescent dye may bind to other molecules present in the drop preclude the authors from drawing quantitative conclusions about the distribution of virions in the residue. More recently, Kong and coworkers [17] used fluorescence microscopy (FM) with GFP-labeled particles of vesicular stomatitis virus to show that these virions form agglomerates, chiefly settling around the innermost part of the rim of dry sessile droplets. In this work, however, the authors do not describe the mechanisms that transport the virions to that location. More importantly, their experimental technique does not allow them to determine the structure of the microenvironment where virions lie, a crucial matter to understand the mechanisms that lead to their slow inactivation. Furthermore, their fluorescence-based technique is only sensitive to agglomerates of fluorescence-labeled virions and therefore unable to detect the presence of virions in other regions where still their concentration may be substantial. More recently, Pan and coworkers from Marr’s lab [5] studied the distribution of labeled Influenza A virions in the dry residue of both model respiratory (a mixture of PBS and Mucin) and human saliva droplets evaporating at low, medium, and high relative humidity (RH). For drops with physiological protein concentration and for saliva they observed that virions predominantly concentrate close to the drop’s rim at low (20%) and medium (50%) humidity. For high (80%) humidity virions spread more uniformly and even concentrate close to the center. Interestingly, these distributions mainly coincide with those of the protein itself.

Although the above mentioned studies shed light on the distribution of virions in the residue of the drop, they do not explain in a quantitative manner the processes that lead to the observed viral and protein distribution. In other words, to the formation of the microenvironment that dictates how virions degrade. Indeed, we believe that understanding the transport of virions, proteins, and salt in the evaporating respiratory drop is essential to develop quantitative viral inactivation models based on first principles. We highlight that the influence of the environmental conditions on which the drop evaporates affects the viral infectivity through their effect on the drop hydrodynamics [2, 10, 5]. With this motivation, in this work we study experimentally and theoretically the transport of virions, protein, and salt in model respiratory sessile droplets.

In contrast to other studies, we use Transmission Electron Microscopy (TEM) to examine the dry residue of model respiratory droplets and unequivocally identify virions. As we show below, this technique allows us to discover features of virion distribution, in particular their structure at the microscale, which is not available to other microscopy technique. To achieve this, we lower the concentrations of non-volatile components in the droplet to a point where we can still observe individual virions without altering significantly the evaporation-driven flow that develops within during the drying process. As experimental viral model, we use the porcine coronavirus transmissible gastroenteritis virus (TGEV) that belongs to the genus alphacoronavirus of the *Coronaviridae* family within the *Nidovirales* order [18]. Like other coronaviruses, TGEV is an enveloped, spherical virus with a single-stranded positive-sense RNA genome and a diameter of approximately 110 nm [19]. Structurally, TGEV closely resembles human coronaviruses of significant public health concern, such as SARS-CoV-2, making it an excellent surrogate for studying coronavirus dynamics and stability in respiratory droplets [14]. Notably, TGEV can be propagated to high titres, it is easy to purify, and can be handled in a BSL-2 laboratory, which simplifies experimental procedures. Since we can purify TGEV with high titres, we are able to observe a large number of virion-laden structures with a reasonably small number of experiments and hence to draw conclusions about the virion distribution. Additionally, we also carry out a theoretical study, validated with dedicated experiments without virions, to elucidate the physical mechanisms that shape the structure and distribution of virions and other solutes observed in the dry drop residue using TEM. We use the conclusions drawn from the numerical solution of our theory to explain and rationalize the observed viral distribution, as well as the microenvironment that surrounds virions in the drop residue.

## 2 Materials and Methods

### 2.1 Culture of cells and viruses

Epithelial swine testis (ST) cells (ATCC CRL-1746), originally described by McClurkin *et al*. [20], were cultured at 37 °C in a humidified atmosphere with 5 % CO_2_. Cells were maintained in Dulbecco’s Modified Eagle Medium (DMEM) supplemented with 10 % fetal bovine serum (FBS), 1 mmol L^−1^ sodium pyruvate, 100 U/mL penicillin, 100 µg mL^−1^ streptomycin, 2 mmol L^−1^ glutamine, and 1 mmol L^−1^ non-essential amino acids (hereafter referred to as growth medium). The TGEV strain PUR-MAD [21] was kindly provided by L. Enjuanes (CNB-CSIC, Madrid, Spain). The virus was propagated in ST cells using growth medium supplemented with 2 % FBS and titrated by plaque assay on ST cells monolayers, as previously described [22, 23].

### 2.2 Virus purification

Highly purified TGEV was prepared as previously described [24]. Briefly, ST cells were infected with TGEV at a multiplicity of infection (MOI) of 1 plaque-forming unit (PFU) per cell. When the cytopathic effect (CPE) was apparent (36 h post-infection), the culture supernatants were harvested and clarified by centrifugation at 4500 *g* for 20 min at 4 °C. The virus was then concentrated by ultracentrifugation at 112000 *g* for 120 min at 4 °C through a 31 % (w/w) sucrose cushion in TEN buffer (10 mM Tris-HCl [pH 7.4], 1 mM EDTA, 1 M NaCl) with 0.2 % Tween 20 and subsequently purified over a continuous 30 % to 42 % sucrose density gradient (112000 *g* for 120 min at 4 °C). Finally, the purified virus was centrifuged at 112000 *g* for 60 min at 4 °C, resuspended in TNE buffer (10 mM Tris-HCl [pH 7.4], 1 mM EDTA, 100 mM NaCl), aliquoted and stored at −80 °C until use. Virus concentration and virus titre were determined by spectrophotometry and plaque assay, respectively. Finally, purity and structural integrity of the viral preparation were confirmed by conventional electron microscopy, which showed that over 99 % of particles retained their morphology, with no detectable vesicles or other contaminants.

### 2.3 Drop compositions used in the TEM experiments

To generate the drops we used artificial saliva, purified TGEV (titre 1.27 × 10^9^ TCDI50) in TNE, and 16 % parafolmaldehyde. The artificial saliva is prepared following the composition used by Vejerano & Marr [11], namely milli-Q water with 9 g L^−1^ of NaCl (Sigma-Aldrich), 3 g L^−1^ of porcine gastric mucin type III (Sigma-Aldrich) and 0.5g L^−1^ of the pulmonar surfactant 1,2-dihex-adecanoyl-sn-glycero-3-phosphocholine. Paraformaldehyde was added to inactivate virions prior to handling outside BSL-2 facilities.

Two saliva-virus solutions were prepared with different levels of dilution (Compositions A and B), as well as blank solutions where the virus was replaced with TNE buffer (Composition Blank A and Composition Blank B):

**Table 1:**
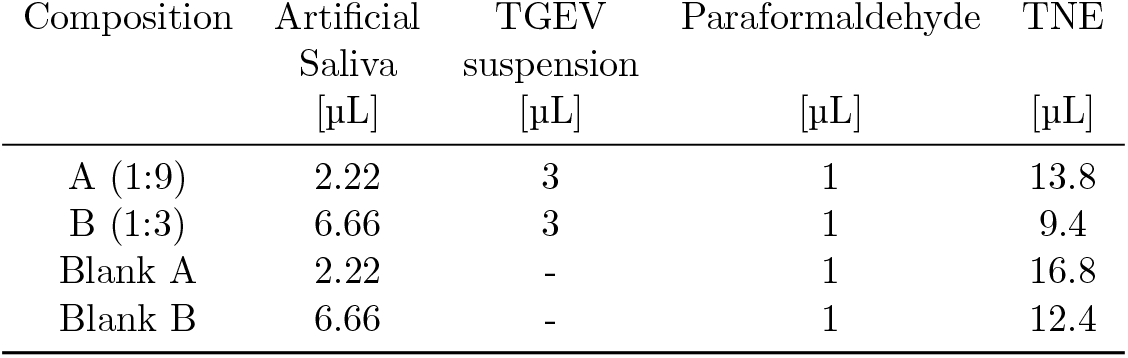
Drop compositions used in the experiments.

Using these compositions, we carried out the following TEM experiments:

**Table 2:** List of TEM experiments.

| Composition | Rel. Humidity (%) | Number of replicates |
| --- | --- | --- |
| A | 40 | 3 |
| A | 70 | 3 |
| B | 40 | 3 |
| Blank A | 40 | 3 |

### 2.4 Transmission Electron microscopy

Droplets of samples 0.3 *±* 0.1 µL were adsorbed onto Formvar carbon-coated 200 mesh nickel grids (Electron Microscopy Sciences, Hatfield, PA, USA). We recorded the drying process of all droplets whose dry residue was subsequently analyzed by TEM (see Fig S1 in section A.1 for details on the drying kinetics). Once we observed the formation of salt crystals and the cessation of movement within the droplet, the TEM grid was collected and stored in a Petri dish. For all experiments, the dry residue was imaged between 30 minutes and 3 hours after complete evaporation. This protocol was applied consistently, and similar features were observed across three independent replicates for each relevant condition. Unstained grids were visualised in a JEOL JEM 1400 Flash electron microscope (JEOL, Tokyo, Japan) operating at 100 kV. No uranyl acetate staining was used to avoid dilution of salt crytals in aqueous solution. Micrographs were taken with a Gatan OneView digital camera (Gatan, California, USA) at various magnifications.

Control experiments without virions (composition blank A, Fig. S7) and those with TNE only (without protein, Fig. S8), including the negative staining procedure after drying, are presented in Supplementary Information Section A.2.1.

### 2.5 Fluorescence Microscopy

To study the spatial distribution of mucin and salt crystals during droplet evaporation, we prepared *V*_0_ = 0.3*±* 0.1 µL droplets of artificial saliva. The droplets were deposited onto *µ*-Dish 35 mm, high glass-bottom dishes (Ibidi, Germany) under ambient conditions. The composition of the droplets was identical to that used in parallel TEM experiments. Evaporation dynamics were monitored using an inverted widefield fluorescence microscope (Leica DMi8) equipped with a CoolLED pE-4000 illumination system and an Orca-Flash 4.0 sCMOS camera (Hamamatsu, Japan). Images were acquired using a 10×/0.32 HC PL FLUO objective. Excitation was performed using a 405 nm LED set to 25 % power. The microscope was equipped with a quad-band fluorescence filter set (QUA), however, only the channel corresponding to excitation at 405 nm and emission collection at 420–480 nm was used in this study. Both mucin and salt crystals emit fluorescence within this spectral range upon 405 nm excitation, allowing simultaneous visualization of their distribution. The exposure time was set to 20 ms per acquisition. Droplet evaporation was imaged every 30 seconds for a total duration of 10 minutes, or until complete evaporation and residual pattern formation were observed. We have carried out a calibration in a previous study [25] following the ideas of Kajiya and co-workers [26], to ensure that the light intensity is proportional to the total amount of protein integrated along the microscope’s optical axis, the vertical direction, from the substrate to the droplet–air interface.

During the final stages of evaporation, salt crystals effloresce, forming dendritic structures (Figure 2c). Although both mucin and salt crystals are visible under fluorescence microscopy (FM) with excitation at 405 nm and emission detection between 420–480 nm, only mucin exhibits intrinsic aut-ofluorescence in this spectral window [27]. The signal observed from salt crystals is therefore primarily due to light scattering from the protein’s emission. Furthermore, mucin molecules may adsorb onto the surfaces of salt crystals during droplet evaporation, resulting in localized fluorescence signals that coincide with the crystal structures.

### 2.6 Drop evaporation and velocity field measurements

To assess the complex fluid flow within the evaporating sessile droplets, we deposited droplets with an initial volume of *V*_0_ = 0.3*±* 0.1 µL onto a micropetri dish coated with a polymeric substrate (Ibidi, Germany). This dish type enables high-resolution imaging and minimizes autofluorescence. Droplets were evaporated inside a humidity-controlled chamber, based on the design by Boulogne [28]. The relative humidity was varied between RH = 30% and 80%. Humid air was generated by recirculating air through a water bottle, while dry air is produced by passing it through a bottle containing calcium sulfate, a highly hygroscopic salt.

To measure the velocity field inside the evaporating droplets, hollow glass spherical tracer particles (Potters Sphericel 110P8) were introduced at a low concentration (0.003 % w/w) to avoid disturbing the flow. Since the flow is experimentally observed to be axisymmetric, we tracked the particle positions within a meridional plane of the droplet. To achieve this, we illuminated the droplet from below using a thin laser sheet focused through a cylindrical lens. Particle tracking velocimetry (PTV) measurements were recorded from the side. Given that the evaporation timescale is significantly longer than the velocity field timescale, we recorded particle motion at 1 fps and the droplet interface at 0.1 fps. To achieve this, we electrically controlled an LED light to switch on every 10 seconds, briefly illuminating the droplet’s interface.

Light refracted by the droplet interface was corrected analytically using the experimentally determined interface shape and ray tracing, following the method of Kwan and co-authors [29]. Although the velocity field remained qualitatively consistent throughout evaporation, refraction effects became significant when the droplet becomes very thin. Therefore, we analyzed particle trajectories only until the contact angle decreasesed to approximately *ε*_*c*_ ∼ 10°.

To study the final stages of evaporation, we complemented side-view PTV with dedicated experiments using General Defocusing Particle Tracking (GDPT) [30, 31]. Fluorescent polystyrene microparticles (1 µm diameter, Microparticles GmbH) were introduced into the droplets and imaged from below using an inverted epifluorescence microscope (Nikon Eclipse Ti2, Nikon, Japan). GDPT relies on optical systems with shallow depth of field, where particle defocusing correlates with depth position. To enhance performance, GDPT is combined with a cylindrical lens that introduces astigmatism, causing spherical particles to appear elliptical. The ellipse shape varies with depth, enabling accurate 3D tracking. Finally, a calibration set of images was used to build a lookup table that maps particle image shapes to depth.

## 3 Virion distribution in the drop residue

To resolve individual virions in the dry residue, as well as the morphology of their surrounding micro-environment, we use TEM. Note that the fluorescence-based techniques used by other authors [17, 5], while very useful for characterizing the spatial distribution and relative abundance of virions within the droplet residue, do not provide access to the microscale organization of virions or the fine structure of the surrounding matrix. The experiments analyzed under the TEM have been performed on evaporating model respiratory droplets containing TGEV at two relative humidity values: RH = 40 % and RH = 70 %. We summarize the main results from the TEM micrographs in Figure 1; however, for further support of our main claims, we encourage the interested reader to consult the supporting information (see section A.2) and the companion open-access dataset [32] (see Section A.2.2 for details on the structure of the repository). Specifically, Figure 1a shows a sketch of a droplet evaporating on a TEM grid. The contact line, which is pinned at all times due to the presence of the protein, is indicated with a dashed cyan line that is also shown in panel (b). This panel presents a global view of the drop residue seen on the TEM grid (black perpendicular bars), where the solid yellow line indicates the microscope field of view. Panels (c) and (d) show two micrographs acquired in the region close to the contact line (highlighted in red in panel (b)), while panels (e) and (f) display the central region of the droplet residue (highlighted in green in panel (b)).

**Figure 1:**
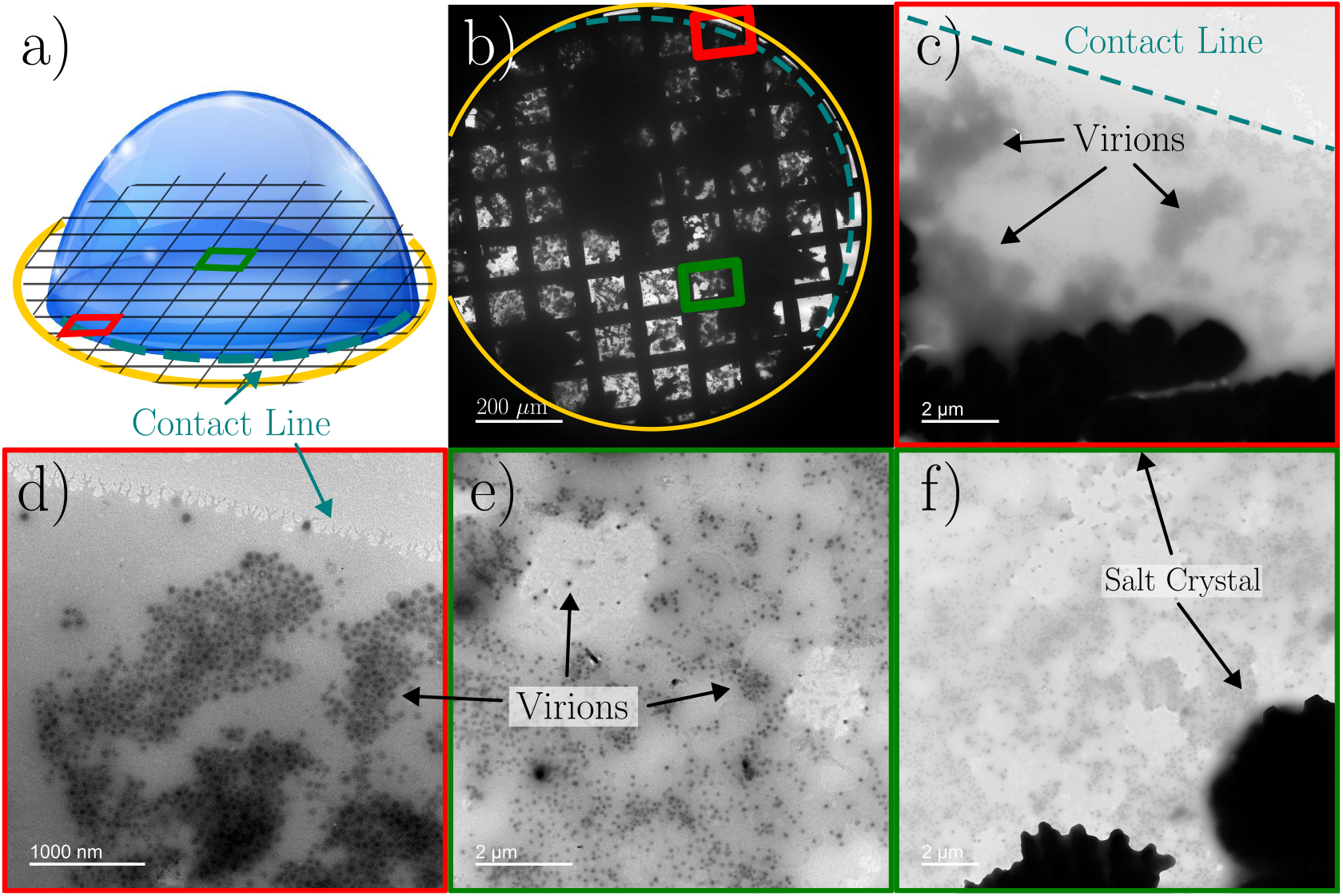
TEM images of different regions of the dry residue of a drop of composition A evaporated at RH = 40 %. (a) Sketch of the geometry, showing the field of view of the microscope (yellow), the contact line (dashed cyan) and two regions, one close to the contact line (red) and another one near to the center (green). (b) Overall TEM view of the whole residue on the TEM grid, indicating the field of view, the contact line and the two selected regions. (c) and (d) Two images inside the red square of (b). We can see a dendritic salt crystal at the bottom of (c). We point at the contact line and the virions, most of them appearing in aggregates. Panels (e) and (f) portrait images belonging to the green square of (b), away from the rim.

At first glance, our TEM images support the findings of previous studies that employed fluorescence microscopy to investigate the overall distribution of virions [17, 5]. Using humidity values between 20% and 50%, these studies reported a significant accumulation of virions in the rim region of the droplet, an observation consistent with our TEM results at RH = 40 % (Fig. 1c,d, Fig. S2 and Ref. [32]). Crucially, as stated above, in contrast to fluorescence microscopy, TEM allows to see individual virions and their surrounding microstructure after evaporation. This allows us to make three observations regarding the way they arrange in the drop residue.

The first observation is that virions are connected forming large but shallow networks, or aggregates, mostly consisting in monolayers (Fig. 1d). This surprising feature is consistently observed under our TEM conditions, as the depth of field used to acquire the micrographs greatly exceeds the size of a single virion, ensuring that no additional virion layers are present in the images, as those shown in Figs. 1d and S2. Aggregates containing regions with multiple layers of virions, beyond what can be resolved by TEM, are also observed, although less frequently (see Fig. S2a, c). At the lowest evaporation rates, or equivalently for the residues obtained at RH = 70 %, viral aggregates are still visible but they tend to be smaller (Fig. S6). Additionally, the distribution of virions is more homogeneous at RH = 70 %, with more virions in the central region of the residue and fewer close to the rim. We also observe in the central region patches where the substrate is clear, indicating an absence of detectable protein (compare TEM images of drops evaporated at 40% and 70% in the companion dataset [32]).

A second observation is that, either isolated or part of an aggregate, virions are mostly found in regions rich in protein (Figs. 1e, 1f, S3). This is what other authors refer to as *co-localization*. This is observed both in dry and humid conditions, and even inside the rim and in the central portion of the residue. This last point is remarkable, since this central portion contains low concentration of protein [5, 25]. In fact, the relatively low protein concentration found in the central region at RH = 40 % leads to areas where the liquid dewets the substrate. These dewetted regions, mostly free of protein, can be seen in Figs. 1e, 1f, Figs. S3a, S3b, and in more images available in [32]. Despite the low protein abundance in the central region of the drop, it is clear from our images that individual viruses are surrounded by a small protein patch (Fig. 1e, f and Figs. S3a, S3b).

A third observation is that a large fraction of the virions segregate from salt crystals, as hypothesized by Morris and coauthors [1]. Due to the high electron density of salt deposits, TEM does not allow imaging through them. Thus, all the viral structures that we can actually see are segregated from salt deposits. Nonetheless, this conclusion is more evident in images where we can see simultaneously viral aggregates and salt crystals as in Figs. 1c, 1f and Fig. S4 (see also Ref. [32]). In these micrographs we see a dendritic formation, characteristic of salt crystals, at the bottom of the image and viral aggregates outside. We emphasize that we see viral structures clearly segregated from salt deposits, but we cannot state whether virions are trapped into, above or below salt formations.

In summary, the TEM micrographs we have shown in this section are representative images of the residue. The same structures have been observed in additional experiments which can be found in the Supporting Information in section A.2 (see Figs. S2, S3, S4). In this section, we have focused on the results obtained for the composition with the smallest amount of protein, since they yield clearer micrographs and thus best convey the message. Nevertheless, it stands to reason that the observations we report here must extend also to more physiological compositions, as the flow pattern and the protein-laden structures in the residue are comparable for composition B, that is closer to the physiological one. For this composition with higher content in protein we observe comparable structure in the dried residue under TEM (Fig S5).

## 4 Transport of solutes inside the drop

Using TEM, we have been able to elucidate the microenvironment where virions lie in the dried drop residue in ways that are not accessible to other microscopy techniques. However, TEM can only be applied to the static residue, once the droplet has totally evaporated, making this technique inadequate to observe in action the transport mechanisms behind the observed residue structure. With the aim of clarifying the mechanisms shaping the final viral distribution, in particular its surprising organization in shallow layers (mostly monolayers), we have carried out two sets of experiments: in the first one we analyze the spatio-temporal distribution of protein (mucin) as the drop evaporates, taking advantage of the natural fluorescence of mucin (FM, Materials and Methods, Fig.2a-c). In the second set, we measure the fluid velocity field inside the drop using two particle tracking velocimetry techniques: a two-dimensional one applied at the drop’s middle plane, allowing for more statistics in the bulk of the droplet, and a three-dimensional one, allowing for better precision on the particle trajectories in the vicinity of the liquid-air interface (PTV, Materials and Methods, Fig.3a). In both sets of experiments, the droplets contained the same composition, without virions, as those used in the TEM experiments. For PTV experiments we introduce polymer-based tracer microparticles. It is important to point out that we do not expect these polymer-based particles to be perfect virion surrogates. Their role here is to act as tracers to quantify the velocity field and the rate at which material from the bulk is captured at the receding droplet interface, which should be practically independent of the particle nature. In fact, to explain where and how virions are deposited the way they are in the residue of the dry drop, we examine the transport of protein since, as we showed above, virions are mostly found in regions that contain protein.

**Figure 2:**
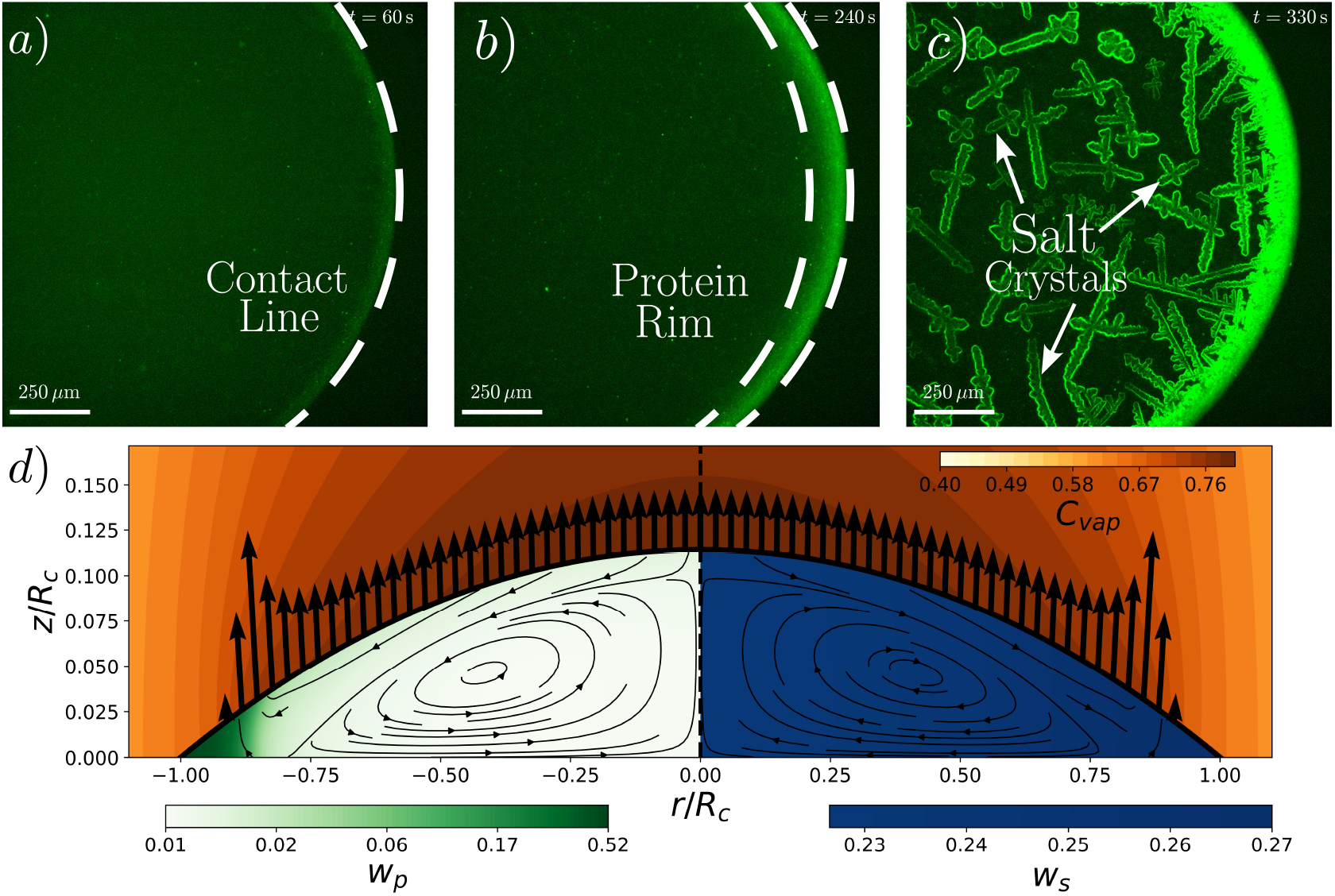
Experimental and numerical study of the formation of the protein rim. (a-c) Three instants during the drop evaporation process (RH = 40%) seen using fluorescence microscopy. From left to right, *t* = 60, 240, 330 s. Protein accumulation near the rim is evident at *t* =240 s. At *t* = 330 s salt crystals have precipitated. The fact that they glow at the same wavelength as the protein suggests that salt crystals are coated by a protein layer. (d) Numerical simulation snapshots of an evaporating model respiratory droplet right before salt saturation is reached at the contact line (RH = 40%). Plotted over the non-dimensional radial (*r/R*_*c*_) and axial coordinated (*z/R*_*c*_) is the vapor concentration, *C*_vap_, represented using a red color scale. On the left side of the droplet, the protein concentration is shown, given by the protein mass fraction, *w*_*p*_, is shown in green, while on the right side, the salt concentration, given by the mass fraction *w*_*s*_, is shown in blue.

The accumulation of protein at the droplet edge over time in the fluorescence microscopy (FM) experiments reveals the existence of a net flow that transports material towards the region close to the rim (Fig. 2(a-c)). The most common source of bulk flow towards the rim in a sessile evaporating drop is the *capillary flow*, a well-known phenomenon firstly identified by Deegan and coworkers [33]. Such an evaporation-driven flow appears naturally due to a mismatch between the evaporative flux and a restricted contact-line motion, typically caused by contact-line pinning [34]. In order to maintain a spherical-cap shape and minimize the surface energy, a capillary liquid flow develops to replenish the contact line region. In the absence of other competing evaporation-driven flows, and for low-diffusivity solutes, the capillary flow concentrates non-volatile materials near the contact line and depletes the central region, leading to the so-called “coffee-ring effect” [33, 34].

However, for drop compositions typical of respiratory fluids not all the material transported to the ring stays there, as it is evident in the FM experiments. In Fig. 2(a-c) we show three FM images taken at the beginning (left), middle (center), and end (right) of the evaporation process. Such a protein distribution is an indication that the capillary flow is not the only transport mechanism present in the evaporation of sessile respiratory droplets, and certainly not the one responsible for the observed viral distribution. In fact, were the capillary flow the only mechanism transporting virions in the drop, then they would all end up at the contact line.

We start by observing the spatial and temporal evolution of the protein. In Fig. 2a and b, before salt precipitates, the fluorescence emission is proportional to the total amount of protein integrated along the microscope optical axis, that is, integrated along the vertical direction from the substrate to the droplet–air interface [26, 25]. In panel 2a, at *t* = 0 s, we see that an incipient protein rim is forming, although a substantial amount of protein appears on the central region. In panel 2b, at *t* = 240 s, the rim is clearly visible and contains a much larger amount of protein than the central region as a consequence of the capillary flow. Interestingly, at the end of the evaporation process, when salt crystals form (panel (c)), there is a large amount of salt in the central region. There is also enough protein to make the salt crystals fluorescent. The results for a drop that evaporates at RH = 70% are qualitatively similar (Supporting Information, Fig. S16).

The complex distribution of both solutes and virions is an indication that multiple evaporation-driven flows might be present within the sessile droplet, as well as other non-hydrodynamic effects, such as the reduction of the evaporation rate near the contact line as a consequence of the large protein concentration reached there [25]. By adding fluorescent flow tracer particles and using appropriate illumination, one can clearly observe the particles following a toroidal-shaped vortex (Fig. 3a). This observation reveals the existence of a recirculatory flow with axial symmetry directed along the free surface from the droplet apex to the rim, and then back from the rim to the droplet center near the solid substrate. This later centerbound flow component contributes to mitigate the accumulation of solutes that would otherwise be generated close to the rim.

**Figure 3:**
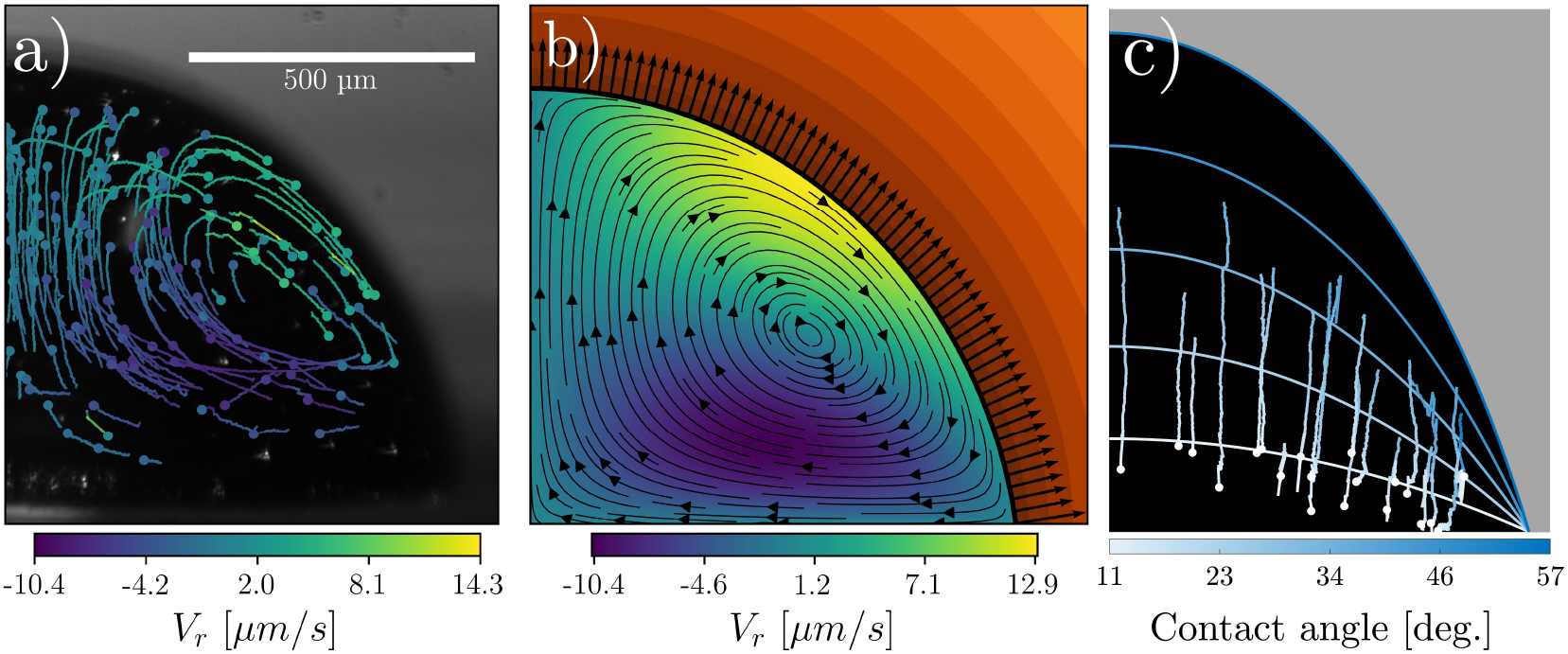
Velocity field inside an evaporating drop. (a) Snapshot of all the particle trajectories detected until *t* = 750 s measured with particle tracking velocimetry for a drop evaporating at RH = 40%. (b) numerical simulation corresponding to the same conditions and time as the experiment on the left panel. (c) Particle trajectories (only those captured at the interface) and interface position detected with a three-dimensional particle tracking technique for a drop evaporating at RH = 40%. We include additional images obtained with this 3D tracking technique in the Supporting Information (Fig. S15).

To explain the physical origin of the recirculation, we have put together a detailed numerical model of the evaporation-induced flow inside the drop. The full description of the model can be found in the Supporting Information (section A.3), but we briefly highlight its main features here. We model the transport of water vapor in the air around the drop as purely diffusive. This leads to an inhomogeneous evaporation rate at the free surface that, thanks to the contact-line pinning, induces a flow inside the drop, slow enough to be dominated by viscous effects [35]. This flow transports non-volatile solutes, mainly salt and protein, and other suspended material like virions. Due to their large concentration, only salt and protein influence the mass transport within the droplet. Therefore, we do not explicitly account for the transport of virions, as their concentration is too low to affect the flow. The mass transport equations both inside and outside the drop, as well as the Stokes equations inside the drop are integrated numerically in time using the finite-element software COMSOL 5.6.

The analysis of the simulations reveals that the high water evaporation rate near the rim makes this region rich in solutes, especially in the case of protein, owing to its lower diffusivity (Fig. 2d). This has several important effects. We start discussing the existence of the recirculation. On the one hand, the high salt concentration near the rim increases the surface tension coefficient. This creates a positive surface tension gradient along the free surface, from the drop apex to the contact line, which translates into a shear stress at the interface (also known as Marangoni stress) which drives a recirculatory flow with the same direction as the one we observe in the PTV experiments, and has been systematically observed in salt-rich evaporating sessile water droplets [36, 37, 38]. Due to their lower concentration, the other solutes present in the drop would generate a surface tension gradient smaller than that associated to the salt. On the other hand, since the liquid density increases with the solute concentration, evaporation induces a larger density near the drop interface, especially close to the rim. As a result, a buoyancy-driven recirculatory flow appears, which also moves fluid in the same direction as the one we observe in the PTV experiments. Since both, interfacial Marangoni stresses and buoyancy-driven forces, contribute to the resultant recirculating flow, a relevant question now is, for the drop sizes and composition used in this work, which one is the most dominant of these two effects on the observed flow? In our numerical simulations we can compute the effect of the density stratification without any free parameter, since the liquid density as a function of the protein and salt concentration can be readily taken from literature [39]. In contrast, it is not possible to calculate the Marangoni stress *a priori*, since this would require knowing how the surface tension depends on the concentration field of surface active material at the droplet’s free surface. This dependency is not easy to characterize experimentally, since tiny amounts of surface contaminants present in the environment, that are impossible to control, have a large impact on surface tension [40, 41, 34]. Consequently, our approach to solve this issue is to find numerically the intensity of the Marangoni flow –the Marangoni number– that best fits our experimentally measured using PTV (more details are given in the Supporting Information, section A). Interestingly, the values found of the Marangoni number are negligibly small, which suggests a negligible contribution of interfacial Marangoni forces to the observed flow pattern. If that is the case, and buoyancy-driven forces are the driving force behind the observed flow, then the direction in which gravity pulls the fluid would be determinant. To corroborate this finding, we conducted experiments with the substrate inclined at 20^*°*^, which clearly breaks the axial symmetry of the flow, confirming the importance of buoyancy-driven forces, see Fig. S14. It is actually not surprising to see that interfacial Marangoni stresses are hindered by contamination or other interfacial effects. There are multiple examples in the literature of such an effect, from pure evaporating sessile water droplets on thermally conducting substrates [42, 34], to rising bubbles in water [43]. In this particular case, the presence of the protein and surfactants *jams* the interface and hinders the appearance of a Marangoni surface stress. The same inhibition of Marangoni due to surface jamming has been observed for droplets containing a solution of albumin [44]. Careful experiments using passive colloidal probes at liquid interfaces have shown that traces amount of surfactant are enough to render the liquid interface incompressible, suppressing the development of surface stresses [45]. To conclude this argument, we would like to insist that the flow pattern but also the velocity values obtained numerically, in the absence of Marangoni stresses, compare remarkably well with experiments without fitting any free parameter (Figs. 3a and 3b).

During the droplet evaporation, a concentrated layer of solute is formed at the drying interface, especially faster at the contact line, where the evaporation flux is stronger. In fact, it can be proven mathematically that the evaporation rate for a droplet of a pure, volatile, liquid is actually infinite at the contact line [33]. To regularize this singularity, as we describe elsewhere [25], we incorporate the effect of the so-called water activity, *χ*, which quantifies how strongly water is retained by the protein matrix versus how available it is for diffusion and/or evaporation. In other words, it represents the ratio –always smaller than one– between the water vapor concentration at our droplet interface (containing proteins, salts and other solutes) and that corresponding to an interface of pure water. The same approach has been followed recently to explain the evaporation insensitivity to relative humidity of drying interfaces of supramolecular solutions [46, 47, 48]. The model reflects the fact that water cannot reach the drop interface for large protein concentrations, since the mutual diffusion coefficient of the water-protein system decays very much for large protein concentrations [49]. In any case, the water activity decreases as the concentration of solutes at the interface increases, leading to a corresponding reduction in the local vapor concentration on the air side. The diffusive flux that drives evaporation, *J*, represented by the black arrows stemming from the drop surface in Figs. 3b and 2, is proportional to the difference between the vapor concentration at the interface and that far away from the drop, thus *J* ∼ *χ* − *H*_*r*_. Consequently, a decrease in water activity reduces the local evaporation rate. For large relative humidity and solute concentration, this effect can even arrest evaporation altogether if the water activity equals the ambient relative humidity [2]. All in all, the reduction in the evaporation rate slows the capillary flow near the rim. This effect is clearly seen in Fig. 2d: the evaporation rate in the protein-rich region close to the rim is negligible compared to that near the center of the drop. We can also see how the maximum evaporation rate is no longer at the contact line, but displaced inwards to a region where the protein concentrations are low enough not to hinder evaporation. This explains why the protein rim, and the virions within, become more distributed than what they would be if all the material was concentrated at the contact line, as would be the case in a suspension of colloids in a pure liquid [33].

An interesting observation, which may have important implications for the viral infectivity [1], is that, at RH = 40 %, whereas protein concentrates mostly near the contact line forming a rim, salt crystals are more homogeneously distributed over the drop residue (Fig. 2c). This is a consequence of diffusion, which opposes to the accumulation of solutes near the contact line. In the case of salt, its diffusivity is still large enough to ensure a rather uniform concentration even at times close to full evaporation. Conversely, the diffusivity of protein is so low that diffusion cannot preclude the formation of the observed rim (Fig. 2c). This conclusion is supported by the values of the Péclet numbers that can be estimated from the measured (and computed) values of the velocity. Using a characteristic velocity *V ≈* 10 µm s^−1^, a drop footprint radius *R*_*c*_ *≈* 0.5 mm, and the following values for the salt and protein diffusivity, *D*_*s*_ = 2 × 10^−9^ m^2^ s^−1^ and *D*_*p*_ = 5 × 10^−12^ m^2^ s^−1^ respectively [50, 51], we have Pe_s_ = *V R*_*c*_*/D*_*s*_ = 2.5 and Pe_p_ = *V R*_*c*_*/D*_*p*_ = 10^3^. These numbers represent the ratio between advective and diffusive transport. Their values confirms that the transport of protein is completely driven by advection (Pe_p_ ≫ 1), while the role of diffusion in the transport of salt cannot be neglected (Pe_s_ ∼ 1) [25].

We verified these transport mechanisms using PIV measurements (Fig S11, S12, S13) and simulations (Fig S18, S17, S19), confirming they are comparable under more physiologically relevant conditions—namely, for droplets of Composition B and artificial saliva (see Section 2.3).

## 5 Discussion

We can draw some conclusions about the transport of virions inside the drop from the examination of the dry drop residue through TEM. We focus on the following observations: (a) virions organize preferentially in oligo- and monolayer aggregates, (b) the aggregates are located everywhere in the residue, but more abundantly in the rim region for RH = 40 %, and (c) almost exclusively surrounded by protein.

It is well known that virions form aggregates both in laboratory cultures and in their natural environment [52]. Among the factors that promote aggregation are the presence of salts [53] and organic materials [54] as well as a free surface [52], all of them elements found in evaporating sessile respiratory droplets. However, the morphology of these aggregates that we report here is more surprising. One would intuitively expect that viral mono- or oligolayers would form more likely with few virions in suspension. Our observations are particularly surprising given the high viral titre that we reach with TGEV (see Materials and Methods). The shallow nature of the observed network suggests that they may have formed at the droplet-air interface, in a mechanism similar to that proposed by Bruning and co-workers [38] for the accumulation of colloidal particles at the liquid-air interface in evaporating NaCl-water droplets. Given the flatness of the observed virion distribution, this mechanism is the most likely to dominate since it has also been previously observed in similar systems.

Such a trapping effect can be clearly observed in our experiments using a three-dimensional particle tracking technique (GDPT [30, 31]) to track fluorescent polymer microparticles suspended in our evaporating sessile droplets containing protein and salts. This technique allows for simultaneous tracking of the particles and the droplet’s free surface, and makes possible to find particles that are immobilized at the liquid-air interface. The results are shown in Fig. 3c, which shows how particles tend to accumulate along the interface especially in the last stages of the evaporation process, when the contact angles are lowest.

Once trapped at the surface, the recirculatory flow transports the particles towards the contact line region, where they aggregate. However, this transport along the interface is eventually arrested, as we discuss in the next paragraph. In the case of virions, it is reasonable to assume that the presence of organic non-volatile material at the interface, in particular proteins, further enhances aggregation. The fact that aggregation takes place to a big extent on a surface endows the aggregates with its two-dimensional nature. If aggregates were formed in the bulk, they would adopt a three-dimensional shape [55].

The capture at the interface, combined with the incompressibility of the liquid-air interface due to the large content in protein, explains the presence of viral aggregates everywhere in the dry residue. In fact, in Fig. 3c we see how captured particles end up following rather vertical trajectories, instead of following the recirculatory flow that dominates the motion in the bulk. This reveals that, after some initial stage, the interface becomes incompressible due to the accumulation of protein. Interestingly, using only salt and a large concentration of colloids, Bruning et al. [38] reported that at high salt and particle concentrations, aggregates are found extending all the way to the center of the drop residue. Recently, a couple of studies [56, 17] have examined the distribution of fluorescently labeled coronaviruses by means of fluorescence microscopy. In both works, the authors found a large fluorescence signal in the central region of the drop, which they identify as virions. In the majority of these studies, the protein rim formation has been explained invoking the classical coffee-ring effect, where all the solutes are advected towards the contact line by the capillary flow [33]. Under the light of our experimental results and quantitative model, such explanation is incomplete and misses many fundamental points.

Another important feature that we can observe thanks to our TEM technique is the presence of viral aggregates close to salt crystal structures. This can be explained as a fluid dynamical process: salt crystals nucleate when the solution becomes slightly supersaturated [57, 58]. At the onset of crystallization, the consumption of salt ions by the growing crystal reduces the saline concentration at its edges down to the saturation concentration, while the saline concentration remains supersaturated far away in the droplet bulk. This concentration gradient creates a flow towards the growing crystal, as reported by Efstratiou *et al*., [59] (we also referred the reader to our movie S6). This flow advects protein, and also virions, towards the growing crystal.

Regardless of the proximity between viral aggregates and salt crystals, it has been proposed that they are protected from the salt’s viricidal effect by either being coated in mucin [1] or forming aggregates [52], whose nucleation and growth is also mediated by the presence of protein at the drop interface. In fact, all the virions that we can see in our experiments –either individuals or members of an aggregate– are located in regions that contain protein (Fig. 1 and Figs. S2 and S3). This observation supports the mechanism proposed by Morris and coworkers [1]: at the end of the evaporation process salt and protein partition and segregate. In this scenario, virions would be found mainly in protein-rich regions, far away from the damaging effect of the salt. To the best of our knowledge, this is the first time that the segregation process underlying this hypothesis has been experimentally verified in a droplet containing real virions, even though evidence that salt and protein segregate during drop evaporation has been known in the literature for years [60, 61].

The observation that virions segregate with protein is in agreement with the conclusion of Wardzala and coworkers [56], who reported that mucin binds to the surface of the coronavirus OC43 particles. It also supports the hypothesis of Alexander et al. [62], who proposed that mucin protects virions from environmental damage, based on infectivity assays performed with varying mucin concentrations in droplets containing mouse hepatitis virus. Interestingly, it has been shown that virions co-localize with proteins for a broad range of viruses (from influenza to other coronaviruses [17, 5]). However, these studies were based on fluorescence microscopy with resolutions much larger than the size of individual viral particles. Thus, they cannot resolve whether co-localization occurs at the level of single virions or larger aggregates. In contrast, our observations confirm that co-localization occurs at the scale of individual virions. Indeed, we do not observe almost any virion that is not surrounded by, at least, a small amount of protein. The same applies to viral aggregates, which are always found in the presence of protein.

In summary, in this work we have contributed to answer the question raised by Wilson Poon and coauthors in their review article: “where are the virions in the respiratory droplets? The answer is poorly known at present” [12]. Moreover, to rationalize the observed viral distribution using TEM, we use a combination of fluorescence imaging, advanced microscopic particle tracking techniques and a precise mathematical modeling of the evaporation-driven transport within the sessile droplet. Our PTV experiments and numerical simulations reveal that the buoyancy-driven recirculatory flow inside the drop, along with the capturing of drop material by the receding interface, explain the observed viral distribution. Understanding the evaporation-driven transport processes inside virus-laden respiratory droplets will allow researchers to clarify crucial open questions regarding disease transmission. In fact, the local microenvironment where virions rest determines the viral stability [10], and this microenvironment is ultimately shaped by the transport mechanisms that we elucidate in this article. However, the advantage of being able to study the microenvironment of viral aggregates in the dry residue with TEM is accompanied by three primary caveats. First, the technique is inherently static, which prevents the study of dynamic virion transport during droplet evaporation. Second, the high resolution required at the nanometer scale precludes efficient mapping of the entire droplet within a reasonable timescale. Finally, the high electron density of salt crystals obscures the internal structure, making it impossible to determine whether virions are embedded within the crystals.

A few years ago, we experienced globally the devastating effects of viral airborne transmission during the COVID-19 pandemic. A natural continuation of this work would be to analyze the viral distribution in more realistic systems, namely airborne droplets made up of real saliva. However extending our experimental methodology to spherical airborne drops is far from trivial. Although there exist some notable experimental studies on the evaporation of such model respiratory drops [16, 8], current techniques do not allow for assessing viral particle distributions within micrometric spherical droplets. Addressing this challenge is essential for advancing our understanding of airborne disease transmission. Last but not least, our work clearly shows the strong heterogeneity of the viral microenvironment. This underscores the need of developing techniques to assess viral viability with enough spatial resolution to distinguish between different regions of the drop residues.

The authors acknowledge financial support from Grant No. PID2023-146809OB-I00 funded by MI-CIU/AEI/10.13039/501100011033 and by ERDF/UE and Grant No. PID2020-114945RB-C21 funded by MCIN/AEI/10.13039/501100011033. We are indebted to Dr. Kevin Roger for his insightful comments about the segregation of components in the dry residue of the drop. We are also very thankful to Prof. Lorenzo Botto for his valuable suggestions on the topic of the formation of viral aggregates on the drop surface.

## References

[1] Dylan H Morris et al. “Mechanistic theory predicts the effects of temperature and humidity on inactivation of SARS-CoV-2 and other enveloped viruses”. In: Elife >10 (2021), e65902.

[2] Carola Seyfert et al. “Stability of respiratory-like droplets under evaporation”. In: Physical review fluids 7.2 (2022), p. 023603.

[3] Aline Schaub et al. “Salt-mediated inactivation of influenza A virus in 1-µl droplets exhibits exponential dependence on NaCl molality”. In: bioRxiv (2023).

[4] Michael D Vahey and Daniel A Fletcher. “Influenza A virus surface proteins are organized to help penetrate host mucus”. In: Elife 8 (2019), e43764.

[5] Jin Pan et al. “Mucin Colocalizes with Influenza Virus and Preserves Infectivity in Deposited Model Respiratory Droplets”. In: Environmental Science & Technology (2025).

[6] Andrea J. French et al. “Environmental Stability of Enveloped Viruses Is Impacted by Initial Volume and Evaporation Kinetics of Droplets”. In: mBio 14 (2023), e03452–22.

[7] Nicole C. Rockey et al. “Seasonal influenza viruses decay more rapidly at intermediate humidity in droplets containing saliva compared to respiratory mucus”. In: Applied and Environmental Microbiology 90 (2024), e02010–23.

[8] Henry P Oswin et al. “The dynamics of SARS-CoV-2 infectivity with changes in aerosol microenvironment”. In: Proceedings of the National Academy of Sciences 119.27 (2022), e2200109119.

[9] Robert Groth et al. “Toward Standardized Aerovirology: A Critical Review of Existing Results and Methodologies”. In: Environmental Science & Technology 58.8 (2024), pp. 3595–3608.

[10] Alexandra K Longest et al. “Review of factors affecting virus inactivation in aerosols and droplets”. In: Journal of the Royal Society Interface 21.215 (2024), p. 20240018.

[11] Eric P Vejerano and Linsey C Marr. “Physico-chemical characteristics of evaporating respiratory fluid droplets”. In: Journal of The Royal Society Interface 15.139 (2018), p. 20170939.

[12] Wilson CK Poon et al. “Soft matter science and the COVID-19 pandemic”. In: Soft matter 16.36 (2020), pp. 8310–8324.

[13] Lakshmi Nathan and Susan Daniel. “Single virion tracking microscopy for the study of virus entry processes in live cells and biomimetic platforms”. In: Physical Virology: Virus Structure and Mechanics (2019), pp. 13–43.

[14] Miguel Cantero et al. “Monitoring SARS-CoV-2 surrogate TGEV individual virions structure survival under harsh physicochemical environments”. In: Cells 11.11 (2022), p. 1759.

[15] Xiaoyun Yang et al. “Immobilization of pseudorabies virus in porcine tracheal respiratory mucus revealed by single particle tracking”. In: PloS one 7.12 (2012), e51054.

[16] Erik Huynh et al. “Evidence for a semisolid phase state of aerosols and droplets relevant to the airborne and surface survival of pathogens”. In: Proceedings of the National Academy of Sciences 119.4 (2022), e2109750119.

[17] Zi-Meng Kong et al. “Virus dynamics and decay in evaporating human saliva droplets on fomites”. In: Environmental science & technology (2022).

[18] Patrick CY Woo et al. “ICTV Virus Taxonomy Profile: Coronaviridae 2023: This article is part of the ICTV Virus Taxonomy Profiles collection.” In: Journal of general virology 104.4 (2023), p. 001843.

[19] Ella Hartenian et al. “The molecular virology of coronaviruses”. In: Journal of Biological Chemistry 295.37 (2020), pp. 12910–12934.

[20] AW McClurkin and James O Norman. “Studies on transmissible gastroenteritis of swine: II. Selected characteristics of a cytopathogenic virus common to five isolates from transmissible gastroenteritis”. In: Canadian Journal of Comparative Medicine and Veterinary Science 30.7 (1966), p. 190.

[21] Carlos M Sánchez et al. “Antigenic homology among coronaviruses related to transmissible gastroenteritis virus”. In: Virology 174.2 (1990), pp. 410–417.

[22] G Jiménez et al. “Critical epitopes in transmissible gastroenteritis virus neutralization”. In: Journal of Virology 60.1 (1986), pp. 131–139.

[23] Carlos M Sánchez et al. “Genetic evolution and tropism of transmissible gastroenteritis coronaviruses”. In: Virology 190.1 (1992), pp. 92–105.

[24] Aitor Nogales et al. “Transmissible gastroenteritis coronavirus RNA-dependent RNA polymerase and nonstructural proteins 2, 3, and 8 are incorporated into viral particles”. In: Journal of virology 86.2 (2012), pp. 1261–1266.

[25] Javier Martínez-Puig et al. “On the role of water activity on the formation of a protein-rich coffee ring in an evaporating multicomponent drop”. In: arXiv preprint arXiv:2509.09315 (2025).

[26] Tadashi Kajiya*, Daisaku Kaneko, and Masao Doi. “Dynamical visualization of “coffee stain phenomenon” in droplets of polymer solution via fluorescent microscopy”. In: Langmuir 24.21 (2008), pp. 12369–12374.

[27] Joseph R Lakowicz. Principles of fluorescence spectroscopy. Springer, 2006.

[28] François Boulogne. “Cheap and versatile humidity regulator for environmentally controlled experiments”. In: The European Physical Journal E 42 (2019), pp. 1–4.

[29] Kwan Hyoung Kang et al. “Quantitative visualization of flow inside an evaporating droplet using the ray tracing method”. In: Measurement Science and Technology 15.6 (2004), p. 1104. doi: 10.1088/0957-0233/15/6/009.

[30] Rune Barnkob, Christian J. Kähler, and Massimiliano Rossi. “General defocusing particle tracking”. In: Lab Chip 15 (17 2015), pp. 3556–3560.

[31] Rune Barnkob and Massimiliano Rossi. “DefocusTracker: A Modular Toolbox for Defocusing-based, Single-Camera, 3D Particle Tracking”. In: Journal of Open Research Software 9.1 (2021), p. 22.

[32] Javier Martínez Puig et al. TEM and Fluorescence Microscopy images of evaporating respiratory droplets containing viral particles. Version V1. 2026. doi: 10.21950/TWRB7H. URL: https://doi.org/10.21950/TWRB7H.

[33] Robert D. Deegan et al. “Capillary flow as the cause of ring stains from dried liquid drops”. In: Nature 389 (1997), pp. 827–829.

[34] Hanneke Gelderblom, Christian Diddens, and Alvaro Marin. “Evaporation-driven liquid flow in sessile droplets”. In: Soft matter 18.45 (2022), pp. 8535–8553.

[35] Yuri O. Popov. “Evaporative deposition patterns: Spatial dimensions of the deposit”. In: Phys. Rev. E 71 (3 2005), p. 036313. doi: 10.1103/PhysRevE.71.036313.

[36] Virginie Soulié et al. “The evaporation behavior of sessile droplets from aqueous saline solutions”. In: Physical Chemistry Chemical Physics 17.34 (2015), pp. 22296–22303.

[37] Alvaro Marin et al. “Solutal Marangoni flow as the cause of ring stains from drying salty colloidal drops”. In: Physical Review Fluids 4 (2019), p. 041601.

[38] Myrthe A Bruning, Laura Loeffen, and Alvaro Marin. “Particle monolayer assembly in evaporating salty colloidal droplets”. In: Physical review fluids 5.8 (2020), p. 083603.

[39] Antonios G. Mikos and Nikolaos A. Peppas. “Measurement of the surface tension of mucin solutions”. In: International Journal of Pharmaceutics 53.1 (1989). doi: 10.1016/0378-5173(89)90354-2.

[40] A Ponce-Torres, EJ Vega, and JM Montanero. “Effects of surface-active impurities on the liquid bridge dynamics”. In: Experiments in Fluids 57.5 (2016), p. 67.

[41] Duarte Rocha et al. “Evaporating sessile droplets: solutal Marangoni effects overwhelm thermal Marangoni flow”. In: Journal of Fluid Mechanics 1013 (2025), A39. doi: 10.1017/jfm.2025.10208.

[42] Hua Hu and Ronald G Larson. “Analysis of the effects of Marangoni stresses on the microflow in an evaporating sessile droplet”. In: Langmuir 21.9 (2005), pp. 3972–3980.

[43] Harishankar Manikantan and Todd M Squires. “Surfactant dynamics: hidden variables controlling fluid flows”. In: Journal of fluid mechanics 892 (2020), P1.

[44] Fan Du, Liyuan Zhang, and Wei Shen. “The internal flow in an evaporating human blood plasma drop”. In: Journal of Colloid and Interface Science 609 (2022), pp. 170–178.

[45] Mehdi Molaei et al. “Interfacial flow around Brownian colloids”. In: Physical Review Letters 126.22 (2021), p. 228003.

[46] Kevin Roger et al. “Controlling water evaporation through self-assembly”. In: Proceedings of the National Academy of Sciences 113.37 (2016), pp. 10275–10280.

[47] Jean-Baptiste Salmon, Frédéric Doumenc, and Béatrice Guerrier. “Humidity-insensitive water evaporation from molecular complex fluids”. In: Phys. Rev. E 96 (3 2017), p. 032612. doi: 10.1103/PhysRevE.96.032612. URL: https://link.aps.org/doi/10.1103/PhysRevE.96.032612.

[48] Max Huisman et al. “Evaporation of Concentrated Polymer Solutions Is Insensitive to Relative Humidity”. In: Phys. Rev. Lett. 131 (24 2023), p. 248102. doi: 10.1103/PhysRevLett.131.248102. URL: https://link.aps.org/doi/10.1103/PhysRevLett.131.248102.

[49] Tania Merhi et al. “Assessing suspension and infectivity times of virus-loaded aerosols involved in airborne transmission”. In: Proceedings of the National Academy of Sciences 119.32 (2022), e2204593119.

[50] Vincenzo Vitagliano and P. Lyons. “Diffusion Coefficients for Aqueous Solutions of Sodium Chloride and Barium Chloride”. In: Journal of the American Chemical Society 78 (Apr. 1956). doi: 10.1021/ja01589a011.

[51] Xingxiang Cao et al. “pH-Dependent Conformational Change of Gastric Mucin Leads to Sol-Gel Transition”. In: Biophysical Journal 76 (1999), pp. 1250–1258.

[52] Charles P Gerba and Walter Q Betancourt. “Viral aggregation: impact on virus behavior in the environment”. In: Environmental science & technology 51.13 (2017), pp. 7318–7325.

[53] Leonardo Gutierrez and Thanh H Nguyen. “Interactions between rotavirus and Suwannee River organic matter: aggregation, deposition, and adhesion force measurement”. In: Environmental science & technology 46.16 (2012), pp. 8705–8713.

[54] Valesca Anschau and Rafael Sanjuán. “Fibrinogen gamma chain promotes aggregation of vesicular stomatitis virus in saliva”. In: Viruses 12.3 (2020), p. 282.

[55] Devendra Pal et al. “Real-time 4D tracking of airborne virus-laden droplets and aerosols”. In: Communications Engineering 2.1 (2023), p. 41.

[56] Casia L Wardzala et al. “Mucins inhibit coronavirus infection in a glycan-dependent manner”. In: ACS Central Science 8.3 (2022), pp. 351–360.

[57] Julie Desarnaud et al. “Metastability Limit for the Nucleation of NaCl Crystals in Confinement”. In: The Journal of Physical Chemistry Letters 5 (2014), pp. 890–895.

[58] Julie Desarnaud et al. “Hopper Growth of Salt Crystals”. In: The Journal of Physical Chemistry Letters 9 (2018), pp. 2961–2966.

[59] Marina Efstratiou, John Christy, and Khellil Sefiane. “Crystallization-Driven Flows within Evaporating Aqueous Saline Droplets”. In: Langmuir 36 (2020), pp. 4995–5002.

[60] J. Filik and N. Stone. “Analysis of human tear fluid by Raman spectroscopy”. In: Analytica Chimica Acta 616.2 (2008), pp. 177–184. ISSN: 0003-2670. doi: 10.1016/j.aca.2008.04.036. URL: https://www.sciencedirect.com/science/article/pii/S0003267008006958.

[61] LV Bel’skaya and EA Sarf. “The use of IR Fourier spectroscopy of saliva for rapid assessment of the level of lipid peroxidation products”. In: Biomedical Chemistry: Research and Methods 2.2 (2019), e00094–e00094.

[62] Robert W Alexander et al. “Mucin transiently sustains coronavirus infectivity through heterogenous changes in phase morphology of evaporating aerosol”. In: Viruses 14.9 (2022), p. 1856.

[63] E Mikhailov et al. “Interaction of aerosol particles composed of protein and saltswith water vapor: hygroscopic growth and microstructural rearrangement”. In: Atmospheric Chemistry and Physics 4.2 (2004), pp. 323–350.

[64] Yana Znamenskaya et al. “Effect of Hydration on Structural and Thermodynamic Properties of Pig Gastric and Bovine Submaxillary Gland Mucins”. In: The Journal of Physical Chemistry B 116 (2012), pp. 5047–5055. doi: 10.1021/jp212495t.

[65] Christian Diddens, Yaxing Li, and Detlef Lohse. “Competing Marangoni and Rayleigh convection in evaporating binary droplets”. In: Journal of Fluid Mechanics 914 (2021). doi: 10.1017/jfm.2020.734.

