## Supporting Information for "Viral transport in evaporating sessile model respiratory droplets"

### A Supporting Information

#### A.1 Evaporation of droplets on TEM grid

We have recorded the evolution of droplet volume throughout the evaporation process (see Figure S1). We observed that, at early times, the volume decreases following the same curve of a pure water droplet; however, at later times, the evaporation slows due to the presence of solutes (salt and protein) (see Figure S1). This behavior is consistent with the experimental and theoretical results of Seyfert et al. (2022) [2]. In Figure S1, we further confirmed that the presence of virions (TGEV) does not affect the drying dynamics.

#### A.2 TEM images residues

In the main text, we present TEM images for the more diluted composition A evaporated at RH = 40%. To further support our main claims, we include additional images of viral aggregates close to the contact line (see Fig. S2). Figure S3 provides further evidence for the co-localization of virions and proteins: the first row shows the raw TEM images, while the second row presents segmented versions of the same images. In these segmented images, protein-rich regions are highlighted in green based on their gray-level contrast with the background, virions are detected and shown in red, and the contact line is manually drawn in blue for the image corresponding to the droplet edge. We also include Figure S4, which shows the morphology of dendritic salt crystals both near the contact line (Fig. S4a) and in the inner region of the residue (Figs. S4b, c).

We have investigated three physiologically relevant conditions: (i) composition A (diluted 1/9) evaporated at RH = 40%, (ii) composition B (diluted 1/3) evaporated at RH = 40%, and (iii) composition A evaporated at RH = 70%. Although the main text primarily focuses on condition (i), the core conclusions of the study remain consistent across all three cases (see Figs. S5 and S6).

Figure S5 shows TEM images of the dry residue from a droplet of composition B evaporated at RH = 40%. The doubled initial protein concentration in composition B makes the contact line region more difficult to resolve clearly by TEM due to the high protein content of that region (see Figs. S5b, c). However, in the inner region of the residue (Figs. S5d, e, f), dendritic salt crystals and viral aggregates are still clearly observed, consistent with the results shown in Fig. 1.

Figure S6 displays TEM images of the dry residue from a droplet of composition A evaporated at RH = 70%. In Fig. S6a, the contrast between the contact line and the inner region is low, indicating a more uniform protein distribution compared to droplets evaporated at RH = 40%. Virions also appear more evenly distributed throughout the residue. Figure S6b, acquired near the contact line, shows a lower concentration of virions than that observed at RH = 40% (cf. Fig. 1 in the main text). Crystallization still occurs at RH = 70%, with virions observed surrounding the crystals.

##### A.2.1 Control experiments

We have performed two types of control experiments:

1. **Blank Control:** Composition Blank A (the same as composition A but without virions) confirms that features identified in other micrographs as virions do not appear in this control (Fig. S7). Moreover, they demonstrate that the morphology of the dried residue—specifically the formation of a protein rim and the presence of dendritic salt crystals—is not affected by the presence of virions.

2. **Negative Staining Control:** A sample containing the same concentration of virions in TNE buffer but without mucin was prepared and subjected to standard post-drying negative staining using uranyl acetate.

This second procedure was not applied to other droplets because the negative staining dissolved the dry residue, significantly altering its morphology and thus preventing direct comparison with the untreated samples presented in the manuscript. However, it served to verify that the $\sim$ 100 nm diameter particles observed in the mucin experiments were indeed virions. The negative staining allowed for the visualization of individual virion spikes (Fig. S8a) and confirmed that the variety of grayscale intensities appearing in compositions A and B (with protein) arises due to the protein itself. In Fig. S8b, the contrast between the virions and the background is clear, while in Fig. S8c, where protein is present, the virions are embedded in the protein gel, displaying a range of grayscales that contrast with the background.

##### A.2.2 Structure of the open-access repository

To further support the claims made in the main text regarding the viral distribution on the dried residue of model respiratory droplets, we have made more than 150 micrographs available in a public, open repository [32]. This ensures that the representative images shown in the main text and Supplementary Information are supported by a substantially broader dataset. To facilitate access to this volume of data, the repository includes a file named `TEM Images TGEV-Model respiratory drop.pdf`. This file consists of a table that, for each micrograph, describes the droplet composition, the ambient relative humidity during evaporation, the drop number, the specific region of the dried residue (G for global view, R for rim, and C for center), and a brief description of the observed features.

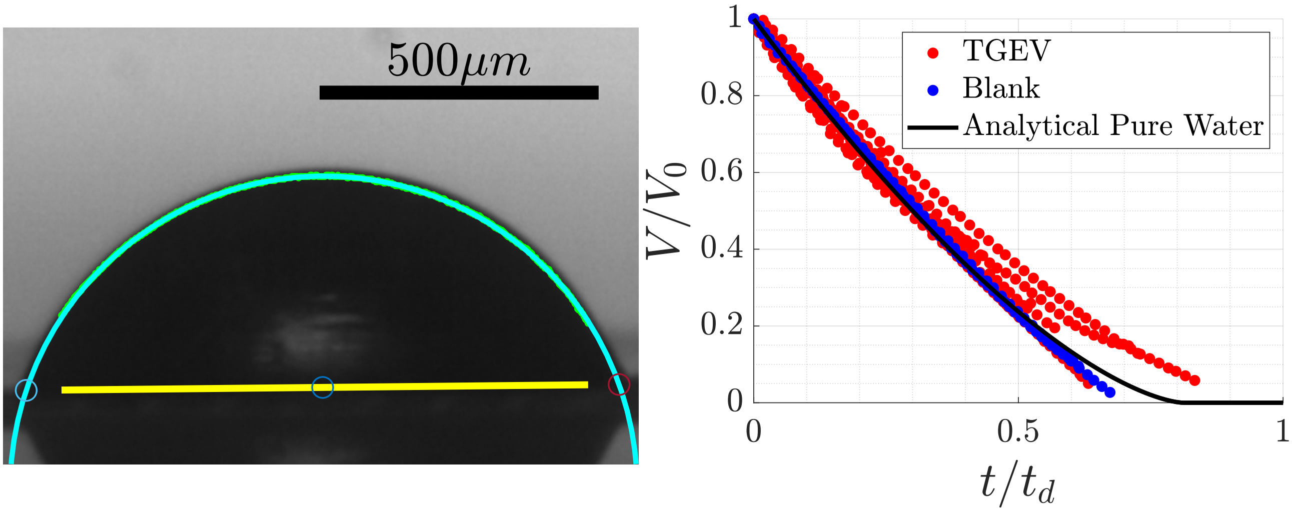


Figure S1: (a) Photo of a sessile drop of composition A evaporating on a TEM grid at $RH = 40\%$. We show the detection of the substrate (yellow line) and the drop surface (cyan line) done by our in-house analysis software. (b) Evolution of the drop volume for several drops evaporating at RH = 40% on a TEM grid. Their evolutions are practically identical, with variations arising from experimental uncertainties. We compare the results with the theoretical curve obtained using the equations in [2].

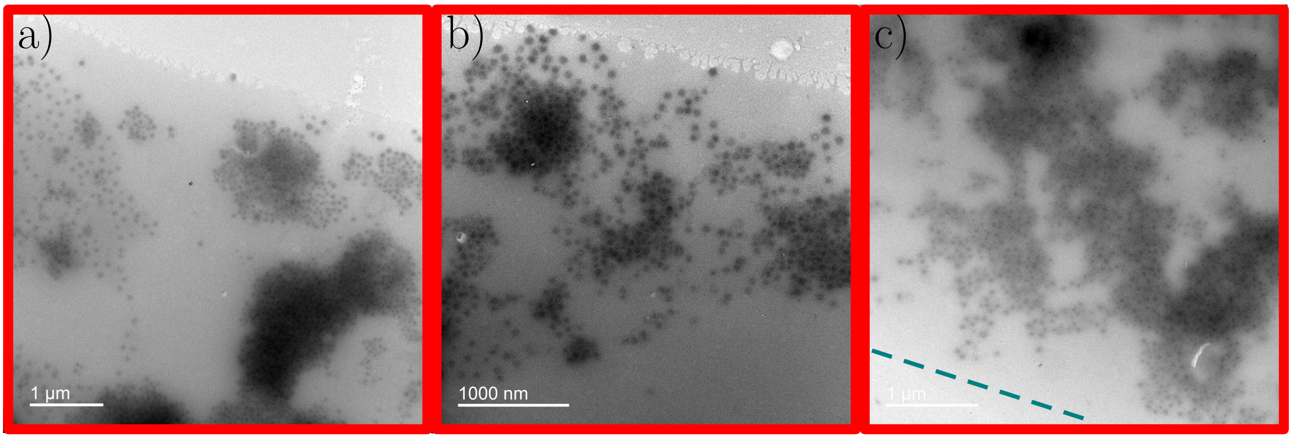


Figure S2: Additional TEM images of the dry residue near the contact line for a droplet of composition A evaporated at RH = 40%. These micrographs highlight the aggregation of virions. The dashed cyan line highlights the contact line in subpanel c).

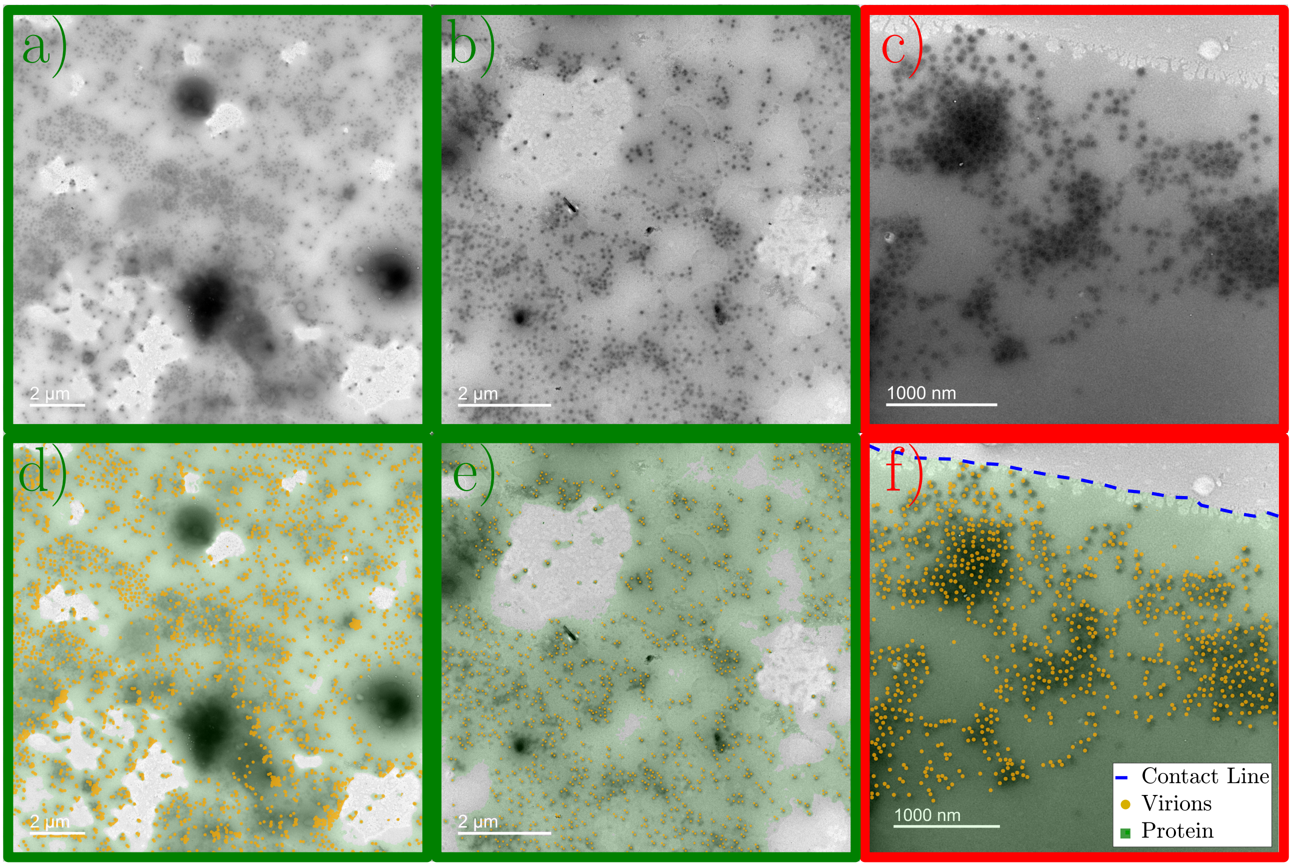


Figure S3: TEM images of virion colocalization with proteins. (a, b) Raw images from the central region of the droplet and (c) the contact line. The second row provides corresponding color-coded interpretations: protein-rich regions are highlighted in green (based on gray-level contrast), virions are indicated in red, and the contact line is delineated in blue.

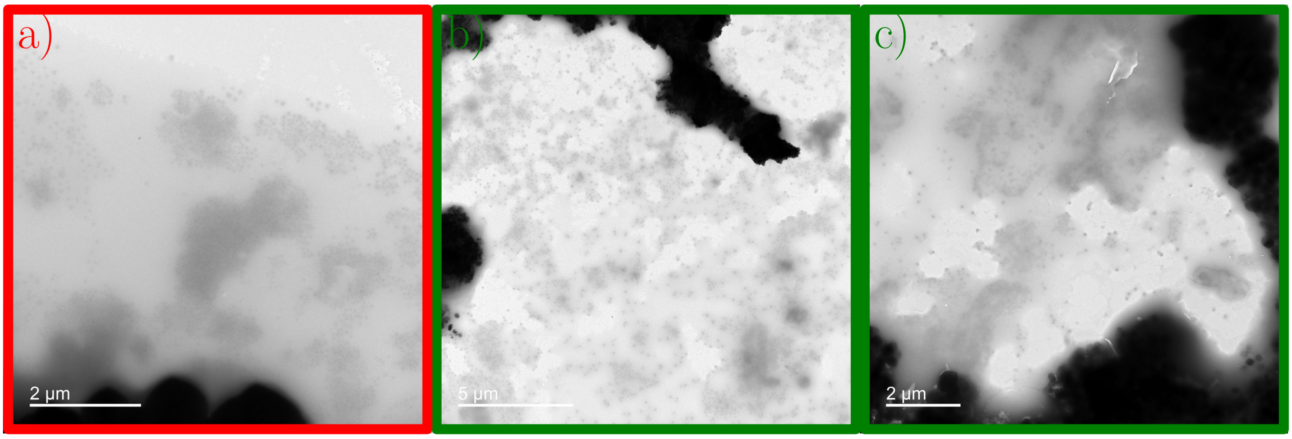


Figure S4: Additional TEM images of the dry residue (a) near the contact line and (b-c) at the center of a droplet (composition A, RH = 40%). These micrographs illustrate the crystal morphology within the dried residue and show that large quantities of virions are segregated from the salt crystals.

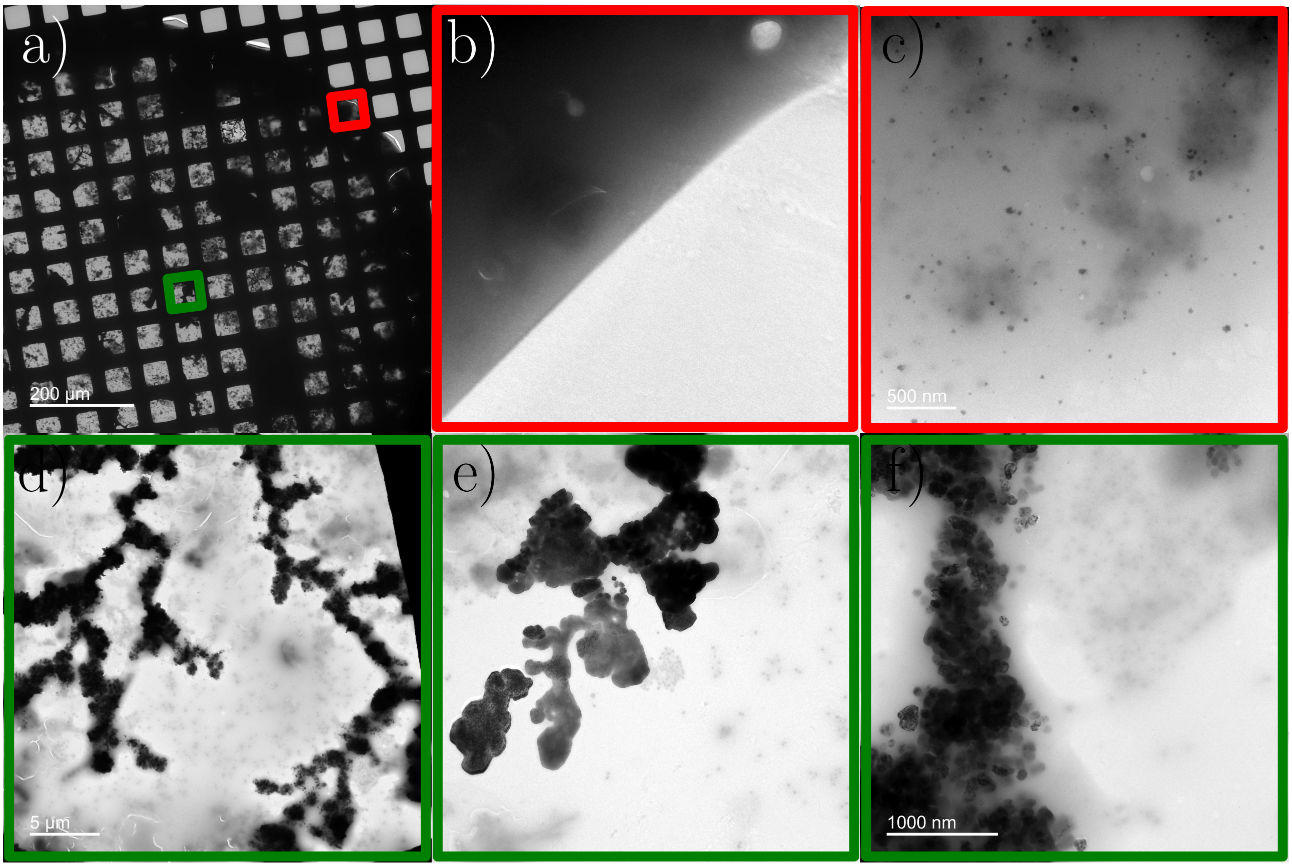


Figure S5: (a) Low magnification image of the dry residue on the TEM grid for a droplet of composition B evaporated at RH = 40%. Panels (b–c) show the region near the contact line, while panels (d–f) correspond to the inner region of the residue.

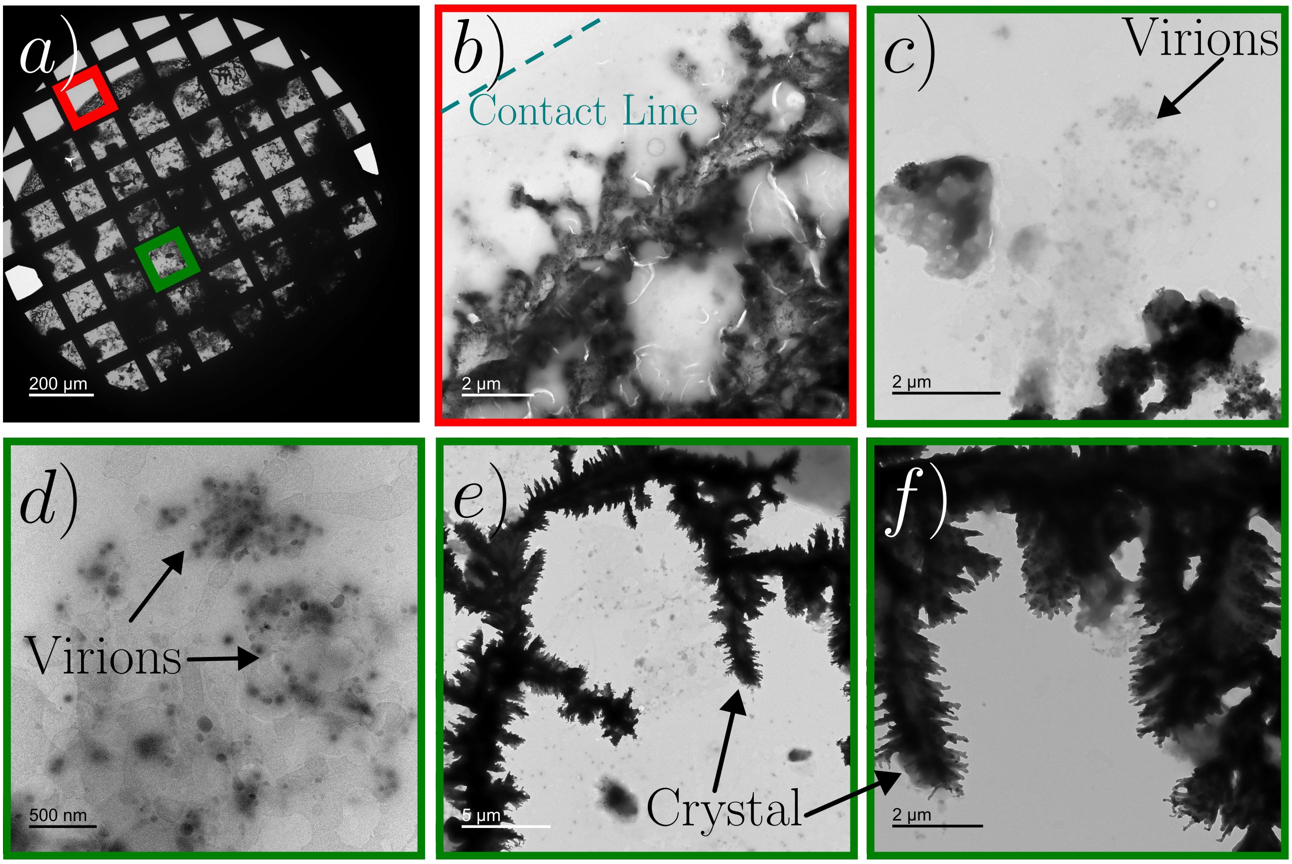


Figure S6: (a) Low magnification image of the dry residue on a TEM grid for a droplet of composition A evaporated at RH = 70%. Panel (b) shows a region near the contact line, while panels (c–f) display the morphology within the inner region of the residue.

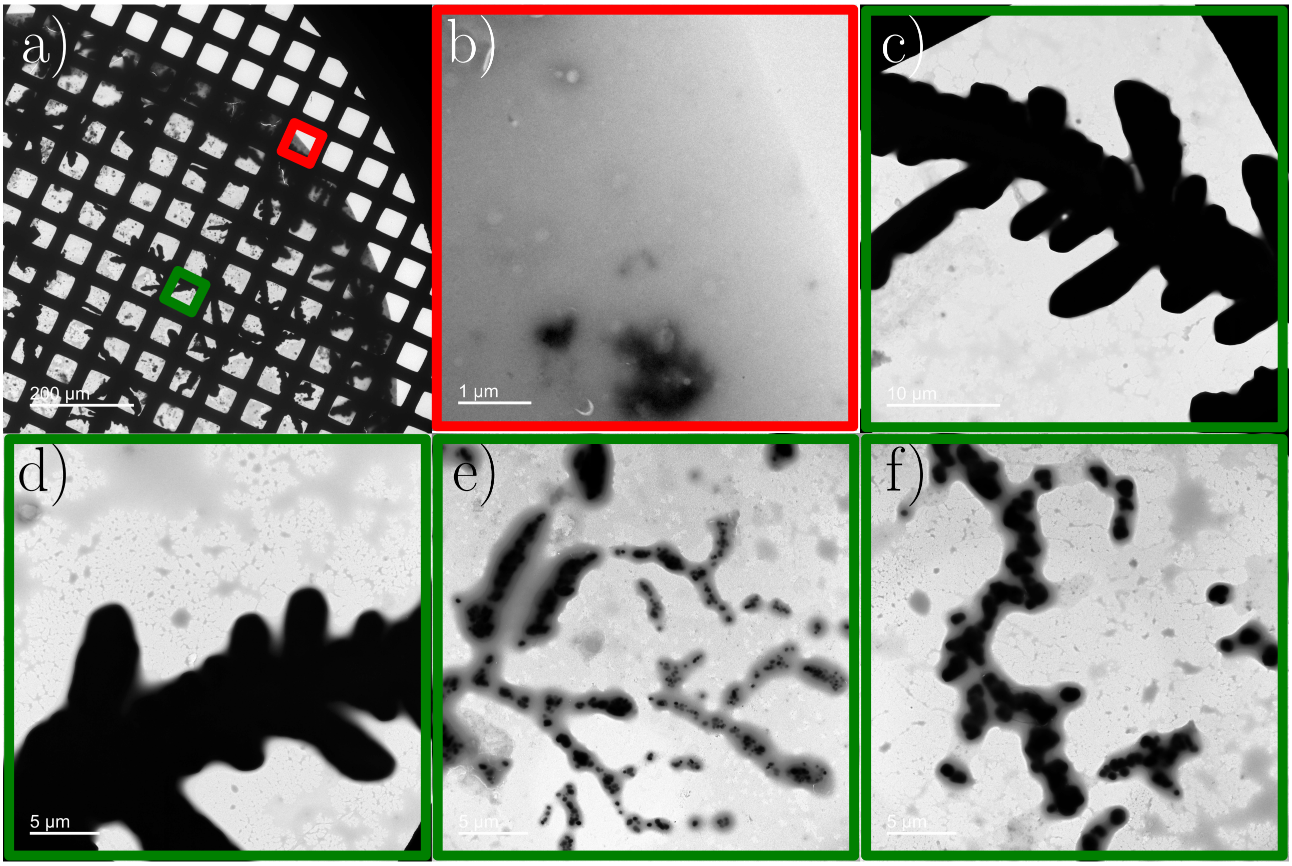


Figure S7: (a) Low magnification image of the dry residue on a TEM grid for a droplet of composition Blank A (without virions) evaporated at RH = 40%. Panel (b) illustrates the region near the contact line, while panels (c–f) display the morphology within the inner region of the residue.

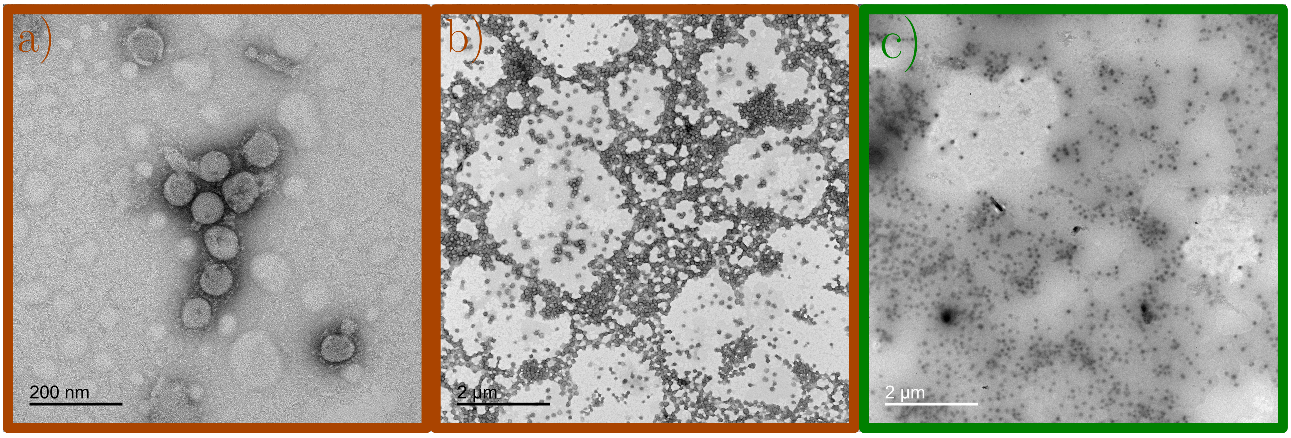


Figure S8: Validation of TGEV virion identification across different media. (a–b) TEM images of virions in TNE buffer (without mucin) with uranyl acetate negative staining: (a) high-magnification image resolving the characteristic viral spikes and (b) a low-magnification image showing distinct contrast against the background. (c) Droplet residue of composition A (with mucin) evaporated at $RH = 40\%$, where virions are embedded within a protein gel, resulting in the varied grayscale intensities and aggregate morphologies observed in the main study.

#### A.3 Hydrodynamical and transport model

The axial symmetry of the problem allows us to formulate the equations in axisymmetric cylindrical coordinates, where $\Omega_G$ and $\Omega_D$ denote the gas and droplet phases, respectively. The domain boundaries are the substrate $\Gamma_{Sub}$, the axis of symmetry $\Gamma_{Sym}$, the far-field $\Gamma_F$, and the gas–droplet interface $\Gamma_I$.

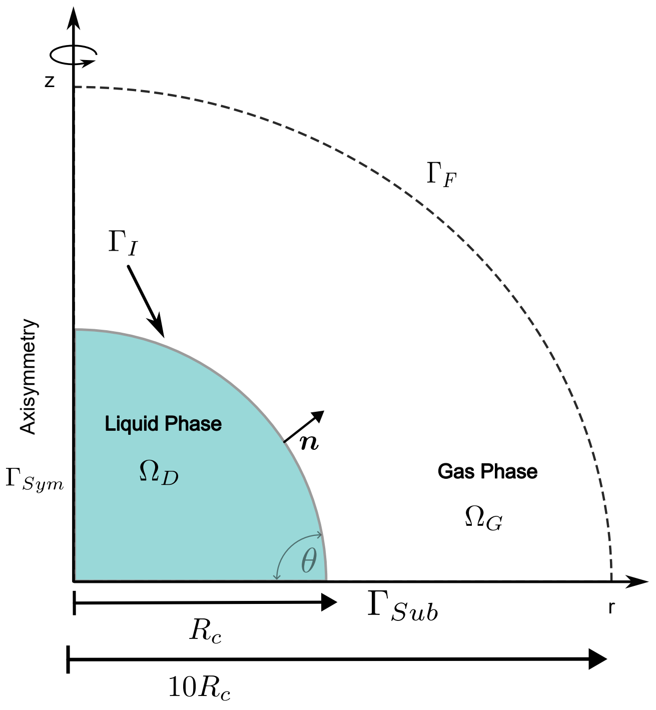


Figure S9: Sketch of the numerical domain.

##### A.3.1 Transport model in the gas phase

In the gas phase we consider the diffusion-driven transport of vapor from the droplet into the air. When the droplet evaporates at a temperature far below the boiling point, and in the absence of forced or strong natural convection, the transport in the gas phase can be considered purely diffusive. Also, the evolution of the vapor concentration field can be treated as quasi-stationary since the diffusive time scale $R_c^2/D_{vap} \sim 10^{-1}$ s, with $R_c$ the radius of the droplet footprint, is several orders of magnitude smaller than the droplet drying time $\rho_0 V_0/(4R_c D_{vap} C_s(1 - H_r)) \sim 10^2$ s. Here, $V_0$ is the initial drop volume, $\rho_0$ the (constant) liquid density, $D_{vap}$ the diffusivity of water vapor in air, $C_s$ the saturation concentration, and $H_r$ the relative humidity.

Under the above assumptions, the equation for the transport of vapor in the air reduces to the Laplacian

$$\nabla^{2*} C^*_{vap} = 0 \text{ at } \Omega_G. \tag{1}$$

We use $*$ to denote the dimensional water vapor concentration and Nabla operator. We describe now the boundary conditions to integrate this equation. At the substrate and the symmetry axis we impose no flux of vapor:

$$\boldsymbol{\nabla}^* C^*_{vap} \cdot \boldsymbol{n} = 0 \text{ at } \Gamma_{Sub} \cup \Gamma_{Sym} \tag{2}$$

At the interphase between the liquid and the gas, thermodynamic equilibrium imposes

$$C^*_{vap} = C_s\, \chi(w_s, w_p) \text{ at } \Gamma_I. \tag{3}$$

Here, $\chi$ is the water activity that depends on the salt and protein mass fraction $w_s$ and $w_p$. To model the water activity we assume $\chi(w_s, w_p) = \chi_s(w_s) + \chi_p(w_p) - 1$ and we interpolate the experimental obtained by Mikhailov and coauthors [63] for $\chi_s$, and by Znamenskaya and coworkers [64] for $\chi_p$.

Far from the droplet, the vapor concentration fulfills

$$C^*_{vap} = C_s\, H_r \text{ at } \Gamma_F \tag{4}$$

We use the vapor saturation concentration $C_s$ to nondimensionalize the vapor concentration $C^*_{vap}$. Doing so, the dimensionless equations for the transport of vapor in air result:

$$\begin{aligned}
\nabla^2 C_{vap} &= 0 \text{ at } \Omega_G && (5)\\
\boldsymbol{\nabla} C_{vap} \cdot \boldsymbol{n} &= 0 \text{ at } \Gamma_{Sub} \cup \Gamma_{Sym} && (6)\\
C_{vap} &= \chi(w_s, w_p) \text{ at } \Gamma_I && (7)\\
C_{vap} &= H_r \text{ at } \Gamma_F. && (8)
\end{aligned}$$

Notice that we omit the $*$ in the dimensionless versions of $\nabla^*$ and $C^*_{vap}$. Then the evaporation rate can be computed as

$$\boldsymbol{J} = -D\lambda \boldsymbol{\nabla} C_{vap} \tag{9}$$

where $\lambda = C_s/\rho_0$ and $D = D_{vap}/D_s$, and $D_s$ is the diffusivity of salt in water. The choice of $J_c = \rho D_s/R_c$ to non-dimensionalize the evaporation rate will be justified later.

##### A.3.2 Transport of salt and protein inside the liquid phase

We will characterize the composition field using the mass fractions, $w_w$, $w_s$, and $w_p$, where the subindexes stand respectively for water, salt and protein. Since these three mass fractions must obey $w_w + w_s + w_p = 1$, we need to write advection-diffusion transport equations for only two of them, namely salt and protein.

We have the two following advection-diffusion transport equations for salt and protein:

$$\partial_{t^*} w_p + \boldsymbol{v^*} \cdot \boldsymbol{\nabla}^* w_p = D_p \nabla^{2^*} w_p \tag{10}$$

$$\partial_{t^*} w_s + \boldsymbol{v^*} \cdot \boldsymbol{\nabla}^* w_s = D_s \nabla^{2^*} w_s. \tag{11}$$

We present now the boundary conditions for these two equations. At the substrate and at the symmetry axis we have a no-flux boundary condition

$$\boldsymbol{\nabla}^* w_p \cdot \boldsymbol{n} = 0 \text{ at } \Gamma_{Sym} \cup \Gamma_{Sub} \tag{12}$$

$$\boldsymbol{\nabla}^* w_s \cdot \boldsymbol{n} = 0 \text{ at } \Gamma_{Sym} \cup \Gamma_{Sub}. \tag{13}$$

At the drop-air interphase the water mass flux must balance the evaporation rate computed by solving the vapor transport at the air side, $J^* = \boldsymbol{J^*} \cdot \boldsymbol{n}$:

$$w_w(\boldsymbol{v}^* - \boldsymbol{v_I}^*) \cdot \boldsymbol{n} - \boldsymbol{J}^*_w \cdot \boldsymbol{n} = \frac{J^*}{\rho_0} \text{ at } \Gamma_I, \tag{14}$$

where $\boldsymbol{v_I}^*$ is the velocity of the drop free surface. Moreover, since the concentration of salt and protein in the gas phase is zero,

$$\begin{aligned}
w_p(\boldsymbol{v}^* - \boldsymbol{v_I}^*) \cdot \boldsymbol{n} - D_p \boldsymbol{\nabla}^* w_p \cdot \boldsymbol{n} &= 0 \text{ at } \Gamma_I && (15)\\
w_s(\boldsymbol{v}^* - \boldsymbol{v_I}^*) \cdot \boldsymbol{n} - D_s \boldsymbol{\nabla}^* w_s \cdot \boldsymbol{n} &= 0 \text{ at } \Gamma_I. && (16)
\end{aligned}$$

We can sum the three boundary conditions and, using $w_w + w_s + w_p = 1$ once more and that the diffusive mass fluxes must sum zero, we arrive to the kinematic boundary condition

$$(\boldsymbol{v^*} - \boldsymbol{v_I^*}) \cdot \boldsymbol{n} = \frac{J^*}{\rho_0} \text{ at } \Gamma_I. \tag{17}$$

This equation will be used to solve for the evaporation-driven normal component of the interphase velocity, $\boldsymbol{v_I}^*$. It also lets us rewrite the boundary conditions for salt and protein mass fractions at the interphase as

$$D_p \boldsymbol{\nabla}^* w_p \cdot \boldsymbol{n} = \frac{J^* w_p}{\rho_0} \text{ at } \Gamma_I \tag{18}$$

$$D_s \boldsymbol{\nabla}^* w_s \cdot \boldsymbol{n} = \frac{J^* w_s}{\rho_0} \text{ at } \Gamma_I. \tag{19}$$

To nondimensionalize our equations we choose

$$\boldsymbol{x} = \frac{\boldsymbol{x^*}}{R_c},\ \boldsymbol{v} = \boldsymbol{v}^* \frac{R_c}{D_s},\ t = t^* \frac{D_s}{R_c^2}. \tag{20}$$

Now we can justify the choice of $J_c = \rho D_s / R_c$ because it leads to a kinematic boundary condition free of parameters,

$$(\boldsymbol{v} - \boldsymbol{v_I}) \cdot \boldsymbol{n} = J. \tag{21}$$

Applying the proposed non-dimensionalization to the rest of equations and boundary conditions we have

$$\partial_t w_p + \boldsymbol{v} \cdot \boldsymbol{\nabla} w_p = \mathcal{D} \nabla^2 w_p \text{ at } \Omega_D \tag{22}$$

$$\partial_t w_s + \boldsymbol{v} \cdot \boldsymbol{\nabla} w_s = \nabla^2 w_s \text{ at } \Omega_D \tag{23}$$

$$\boldsymbol{\nabla} w_p \cdot \boldsymbol{n} = 0 \text{ at } \Gamma_{Sym} \cup \Gamma_{Sub} \tag{24}$$

$$\boldsymbol{\nabla} w_s \cdot \boldsymbol{n} = 0 \text{ at } \Gamma_{Sym} \cup \Gamma_{Sub} \tag{25}$$

$$\boldsymbol{\nabla} w_p \cdot \boldsymbol{n} = \frac{J\, w_p}{\mathcal{D}} \text{ at } \Gamma_I \tag{26}$$

$$\boldsymbol{\nabla} w_s \cdot \boldsymbol{n} = J w_s \text{ at } \Gamma_I. \tag{27}$$

We have introduced the diffusivity ratio $\mathcal{D} = D_p / D_s$.

##### A.3.3 Hydrodynamic equations

In what follows, we will denote the composition-dependent liquid density by $\rho^*$. The evolution of the density is given by the continuity equation

$$\partial_{t^*} \rho^* + \boldsymbol{\nabla^*} \cdot (\rho^* \boldsymbol{v^*}) = 0 \tag{28}$$

Until this moment we have considered the density as a constant, $\rho_0$ but, in fact, the density depends on the composition. That is, on the mass fractions $w_s$ and $w_p$. For dilute solutions, the density can be modeled as a linear function of both mass fractions [39]:

$$\rho^* = \rho_0 + \Delta\rho^* = \rho_0 + \rho_s w_s + \rho_p w_p. \tag{29}$$

However, since $\Delta\rho^*/\rho_0 \ll 1$ only the term $\rho_0$ was retained in all previous equations. An exception is made for the gravity term in the momentum equation, where density variations are maintained. This treatment corresponds to the solutal Boussinesq approximation. To justify neglecting density variations elsewhere, the continuity equation was nondimensionalized usnig the same scaling for time, length and velocity as described, yielding:

$$\partial_t(\rho_0 + \rho_s w_s + \rho_p w_p) + \boldsymbol{\nabla} \cdot ((\rho_0 + \rho_s w_s + \rho_p w_p)\boldsymbol{v}) = 0 \tag{30}$$

Dividing by $\rho_0$ we have that

$$\partial_t \left(1 + \frac{\rho_s w_s + \rho_p w_p}{\rho_0}\right) + \boldsymbol{\nabla} \cdot \left(\left(1 + \frac{\rho_s w_s + \rho_p w_p}{\rho_0}\right) \boldsymbol{v}\right) = 0 \tag{31}$$

Since $(\rho_s w_s + \rho_p w_p)/\rho_0 \ll 1$ during the whole evaporation,

$$\boldsymbol{\nabla} \cdot \boldsymbol{v} = 0. \tag{32}$$

We have arrived at the continuity equation of an incompressible flow which is typical of the Boussinesq approximation.

The momentum equation for an incompressible fluid reads

$$\rho^*(\partial_t \boldsymbol{v^*} + \boldsymbol{v^*} \cdot \boldsymbol{\nabla^*} \boldsymbol{v^*}) = -\boldsymbol{\nabla^*} p^* + \boldsymbol{\nabla^*} \cdot (\mu^*(\boldsymbol{\nabla v^*} + \boldsymbol{\nabla v^{*T}})) - \rho^* g \boldsymbol{e_z}. \tag{33}$$

Our non-dimensionalization, consistent with the one already used in the previous subsections, will be

$$t = \frac{D_s}{R_c^2} t^*, \; \boldsymbol{x} = \frac{\boldsymbol{x^*}}{R_c}, \; \boldsymbol{v} = \boldsymbol{v}^* \frac{R_c}{D_s}, \; \mu = \frac{\mu^*}{\mu_w}, \; P = \frac{R_c^2}{\mu_w D_s}(p^* + \rho_0 g z^*) \tag{34}$$

where $\mu_w$ is the viscosity of pure water and $P$ is the dimensionless reduced pressure. The characteristic pressure $\Delta p_c = \mu_w D_s / R_c^2$ has been chosen because the pressure gradient should be of the same order than the viscous stresses. Applying the non-dimensionalization we have

$$\begin{aligned} \frac{(\rho_0 + \rho_s w_s + \rho_p w_p) D_s}{\mu_w} (\partial_t \boldsymbol{v} + \boldsymbol{v} \cdot \boldsymbol{\nabla v}) = \\ -\boldsymbol{\nabla} P + \nabla \cdot (\mu(\boldsymbol{\nabla v} + \boldsymbol{\nabla v}^T)) - \frac{R_c^3 g}{\mu_w D_s}(\rho_s w_s + \rho_p w_p) g \boldsymbol{e_z}. \end{aligned} \tag{35}$$

Applying again $\Delta\rho^*/\rho_0 \ll 1$ we have

$$Sc^{-1}(\partial_t \boldsymbol{v} + \boldsymbol{v} \cdot \boldsymbol{\nabla v}) = -\boldsymbol{\nabla} P + \boldsymbol{\nabla} \cdot (\mu(\boldsymbol{\nabla v} + \boldsymbol{\nabla v}^T)) - Ra\, \rho\, \boldsymbol{e_z}, \tag{36}$$

where $Sc = \mu_w / D_s \rho_0$ is the Schmidt number, $Ra = R_c^3 g \rho_s / \mu_w D_s$ the Rayleigh number, and $\rho = (w_s + \rho_p/\rho_s\, w_p)$ a dimensionless density variation. For our problem, $Sc > 10^3$, but we can only neglect inertia if the Reynolds number $Re = \rho_0 v_c R_c / \mu_w$ is also small, which is the case in our experiments where $v_c \approx 10\ \mu m/s$ and thus $Re \approx 10^{-2}$. In contrast with this $Ra \sim 10^6$ so we should keep buoyancy forces in the equations. This is the reason why we have applied the solutal Boussinesq approximation: changes in density are only relevant in the buoyancy forces, which only appear in the momentum equation. Finally, the dimensionless momentum equation reads

$$\boldsymbol{\nabla} \cdot \tau - Ra\, \rho\, \boldsymbol{e_z} = 0, \tag{37}$$

where $\tau = -P\mathcal{I} + \mu(\boldsymbol{\nabla v} + \boldsymbol{\nabla v}^T)$ is the dimensionless stress tensor defined using the reduced pressure $P$.

At the symmetry axis we impose symmetry conditions

$$v_r = 0, \; \frac{\partial v_z}{\partial r} = 0 \text{ at } \Gamma_{Sym}. \tag{38}$$

At the substrate we have no slip boundary condition

$$\boldsymbol{v} = 0 \text{ at } \Gamma_{Sub}. \tag{39}$$

At the interphase we have the kinematic boundary condition already derived in the previous subsection which in dimensionless form results

$$(\boldsymbol{v}-\boldsymbol{v_I})\cdot\boldsymbol{n}=J,\ \text{at } \Gamma_I. \tag{40}$$

This equation is needed to compute the interphase as part of our solution. However, this condition only gives the normal component of the interphase velocity, so another equation is needed that includes the tangential velocity component. This is going to be the stress condition

$$-\tau^*\cdot\boldsymbol{n}=\gamma^*(\boldsymbol{\nabla}^*\cdot\boldsymbol{n})\boldsymbol{n}-\boldsymbol{\nabla_t}^*\gamma^*+\rho_0 g z^*\boldsymbol{n}. \tag{41}$$

We assume that surface tension depends linearly on the concentration of salt $\gamma^*=\gamma_0+\gamma_s w_s$. Then in dimensionless form the stress condition is

$$-\tau\cdot\boldsymbol{n}=Ca^{-1}((1+CaMaw_s)\boldsymbol{\nabla}\cdot\boldsymbol{n}+Bo\,z)\boldsymbol{n}-Ma\boldsymbol{\nabla_t}w_s. \tag{42}$$

where $Ca=\mu_w D_s/\gamma_0 R_c$ is the capillary number, $Ma=\gamma_s R_c/\mu_w D_s$ the Marangoni number, and $Bo=\rho g R_c^2/\gamma_0$ the Bond number.

To summarize, the hydrodynamic equations in the droplet are

$$\boldsymbol{\nabla}\cdot\boldsymbol{v}=0,\ \text{at } \Omega_D. \tag{43}$$

$$\boldsymbol{\nabla}\cdot\tau-Ra\,\rho\,\boldsymbol{e_z}=0,\ \text{at } \Omega_D \tag{44}$$

$$v_r=0,\ \frac{\partial v_z}{\partial r}=0 \text{ at } \Gamma_{Sym} \tag{45}$$

$$\boldsymbol{v}=0,\ \text{at } \Gamma_{Sub} \tag{46}$$

$$(\boldsymbol{v}-\boldsymbol{v_I})\cdot\boldsymbol{n}=J,\ \text{at } \Gamma_I \tag{47}$$

$$-\tau\cdot\boldsymbol{n}=Ca^{-1}((1+CaMaw_s)\boldsymbol{\nabla}\cdot\boldsymbol{n}+Boz)\boldsymbol{n}-Ma\boldsymbol{\nabla_t}w_s \text{ at } \Gamma_I. \tag{48}$$

A sketch of the model can be seen in Fig. S10.

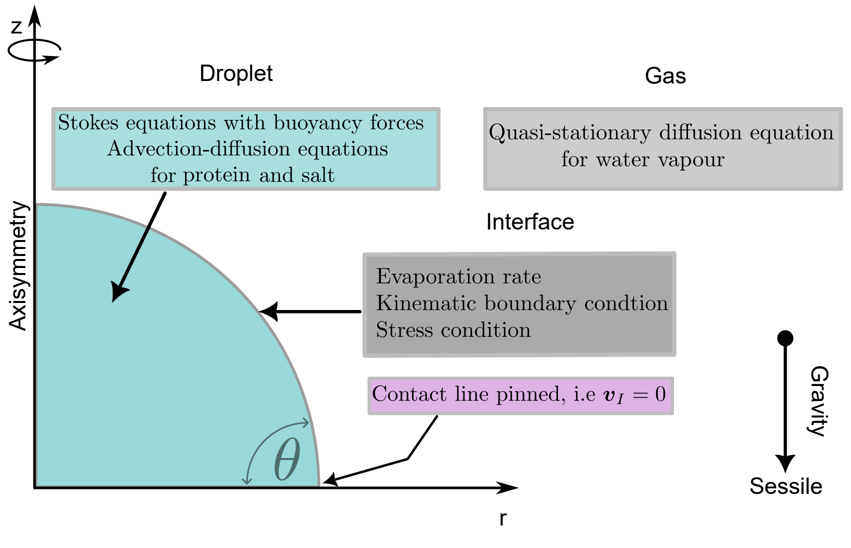


Figure S10: Sketch of the model, detailing which equations are applied in the different domains and boundaries.

This model accounts for the complexity of a respiratory droplet but has been kept as general as possible. Indeed it can be useful to study other ternary mixture droplets, where the solutal Boussinesq approximation can be applied $\Delta\rho^*/\rho^*\ll 1$ and inertia can be neglected. This happens in a wide variety of applications [65]. The particularities of the system can be introduced via the dimensionless viscosity and density, and the dimensionless numbers. As explained in the main text, section "Transport of solutes in the drop", given the negligible Marangoni number derived from our PTV measurements, we set $Ma = 0$ in the simulations.

###### A.3.4 Numerical scheme

To solve the model, we rewrite all dimensionless equations in weak form for implementation in COMSOL Multiphysics using the Mathematics Module. The incompressible Stokes flow with buoyancy forces is spatially discretized using Taylor–Hood triangular elements for velocity and pressure. Mesh deformation due to interfacial motion is handled with the Arbitrary Lagrangian–Eulerian (ALE) method. Time integration is performed using a variable-step, variable order backward differentiation formula (BDF) method of order 2 to 4, with adaptive step size to meet a relative tolerance of $10^{-4}$.

##### A.4 Experimental Velocity Field

###### A.4.1 PTV: Side View

In the main text, we present particle tracking results for composition Blank A at RH = 40%. To complement these results, we include experiments for compositions Blank B (see Fig. S11) and Blank A (see Fig. S12) at the same relative humidity. For both compositions, tracer particles were added as described in Materials and Methods. The higher protein content in these compositions reduces particle visibility, resulting in fewer detected particles and shorter trajectories. Despite this, a recirculatory flow is still observed, with characteristic velocities identical to those in composition Blank A.

Additional particle tracking velocimetry (PTV) experiments were performed at RH = 70% (see Fig. S13). In this case, the recirculatory flow persists but with lower velocities compared to the RH = 40% condition. This is consistent with the hydrodynamic framework presented in the main text, and studied in more detail in Ref. [25]. The slower evaporation rate found at higher humidity translates into slower velocities inside the drop. Also, the distribution of solutes is more uniform, which smooths the density gradient that drives the recirculatory flow.

Finally, to verify that natural convection is the primary mechanism driving the recirculation, we conducted additional experiments in which the droplet was tilted by 20° relative to the direction of gravity. This tilt clearly breaks the flow symmetry (see Fig. S14), and the recirculation cells become approximately separated by the vertical (the direction of gravity), further supporting that the flow is buoyancy-driven.

###### A.4.2 GDPT: Particle capture at the interface

We show here more experiments showing particle trajectories obtained with the GDPT technique. In Fig. S15 we show the particle trajectories obtained during a first stage of the drop evaporation process. For times $t \leq 63s$, the interface has not receded enough to capture a large amount of particles. Thus, they remain in the bulk. If the same analysis is repeated at later times, we see that several tracer particles have been captured by the receding interface. Remarkably, particles trapped at the surface do not follow the recirculatory flow observed in the bulk. This reveals that the interface is jammed with protein and other solutes.

##### A.5 Fluorescence Experiments

We conducted fluorescence imaging experiments on droplets evaporated at RH = 70% (see Fig. S16). Notably, within the timescale of the experiment, no crystallization was observed, in contrast to droplets evaporated at RH = 40%. Additionally, the protein rim is significantly wider under these conditions. The contrast between the center of the droplet and the contact line is also less pronounced at RH = 70%, indicating a lower peak protein concentration within the rim compared to the RH = 40% case. These observations are predicted by the numerical simulations presented in this work.

#### A.6 Additional snapshots of simulations for different conditions

To verify that the flow pattern is similar for both compositions used on our experiments (A and B) and, more importantly, also compared to a droplet that only contains artificial saliva, we carried different simulations. In Fig. S17 we show that the flow pattern of an evaporating droplet of artificial saliva is also recirculatory. Importantly, the order of magnitude of the velocity is comparable to compositions Blank A and Blank B. For completeness, in Fig. S18 we show a snapshot of a simulation of the evaporation of a droplet of composition Blank B at RH = 40%. The simulation snapshots are acquired at a time when the droplet height is comparable.

Finally, we include in Fig. S19 a snapshot of a simulation of an evaporating artificial saliva droplet at RH = 70%. At this relative humidity the evaporation is slower, allowing the solutes more time to diffuse in the vertical direction. As a result, smoother vertical gradients in solute concentration, and consequently in density, are observed. The reduced strength of natural convection accounts for the lower order of magnitude of the velocity field observed at RH = 70%.

#### A.7 Movies

**Movie S1** - Experimental video of PTV on a droplet of composition E, evaporated at Hr=40%.
**Movie S2** - Experimental video of the bottom view used to reconstruct the 3D velocity field thanks to the GDPT technique. Notice that the ellipsoidal shape of the contact line is due to the use of a cylindrical lense to sharp how particles defocused for different optical axis positions.
**Movie S3** - Trajectories obtained with GDPT from Movie S2. The droplet has composition Blank B and was evaporated at Hr=40%.
**Movie S4** - Video of a simulation corresponding to the experiment shown in Fig. 1 on the main text.
**Movie S5** - Video of a fluorescent experiment on a droplet of composition Blank B evaporated at Hr=40%.
**Movie S6** - Evidence of fast flows directed towards salt crystals while they are growing.
