## Supporting Information - Figs. S11-S19 for "Viral transport in evaporating sessile model respiratory droplets"

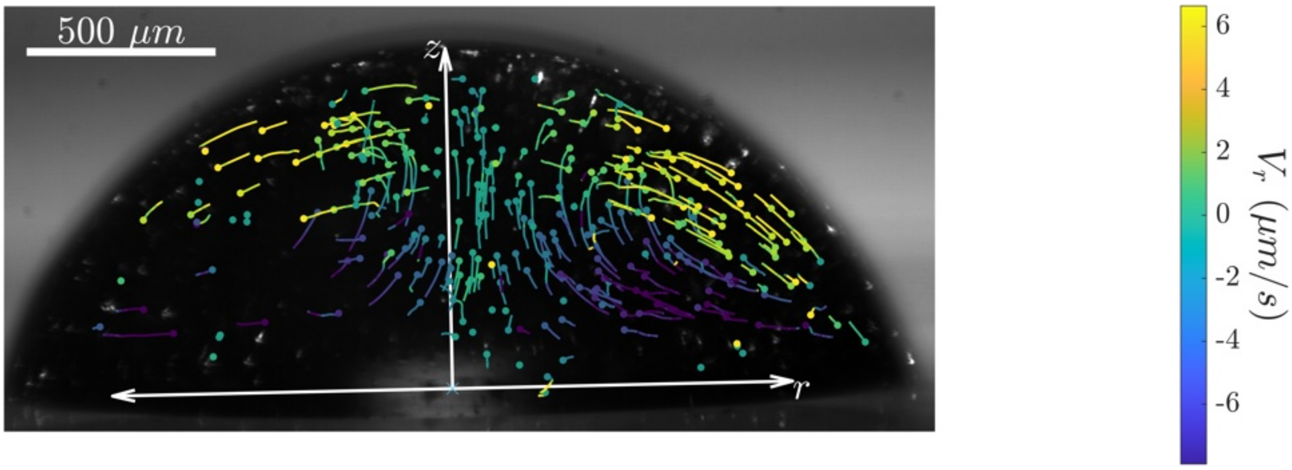


Figure S11: Particle trajectories obtained using PTV on a droplet of composition Blank B evaporated at RH = 40%. The colors represent instantaneous radial velocity.

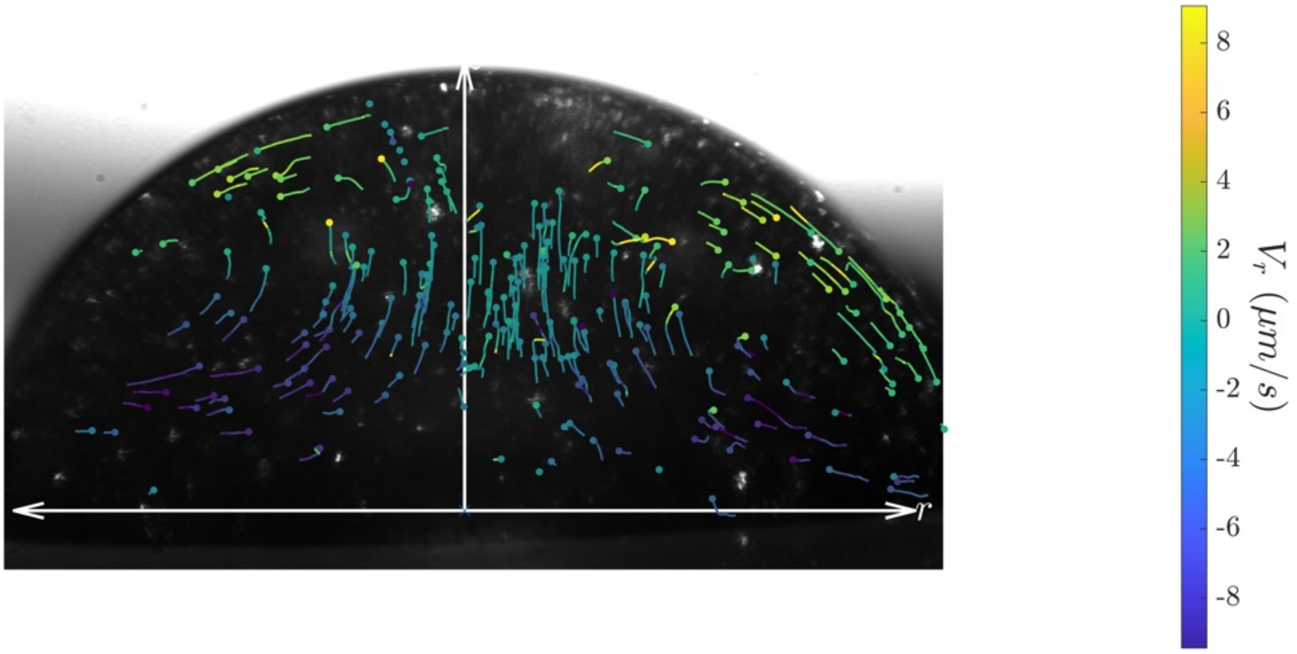


Figure S12: Particle trajectories obtained using PTV on a droplet of composition Blank A evaporated at RH = 40%. The colors represent instantaneous radial velocity.

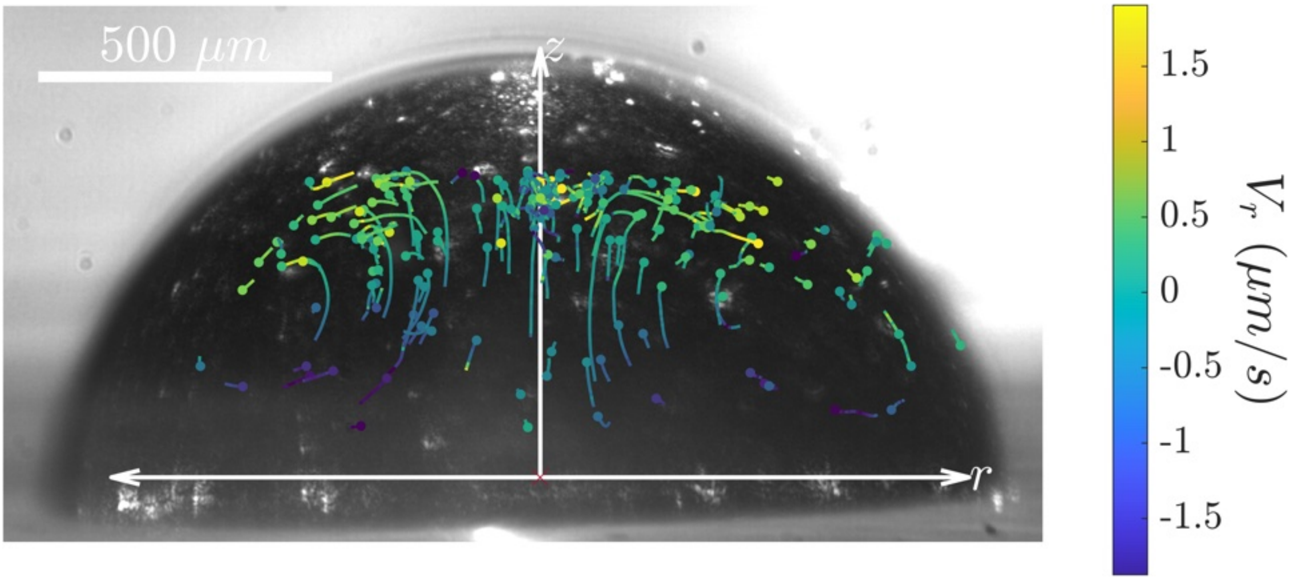


Figure S13: Particle trajectories obtained using PTV on a droplet of composition Blank A evaporated at RH = 70%. The colors represent instantaneous radial velocity.

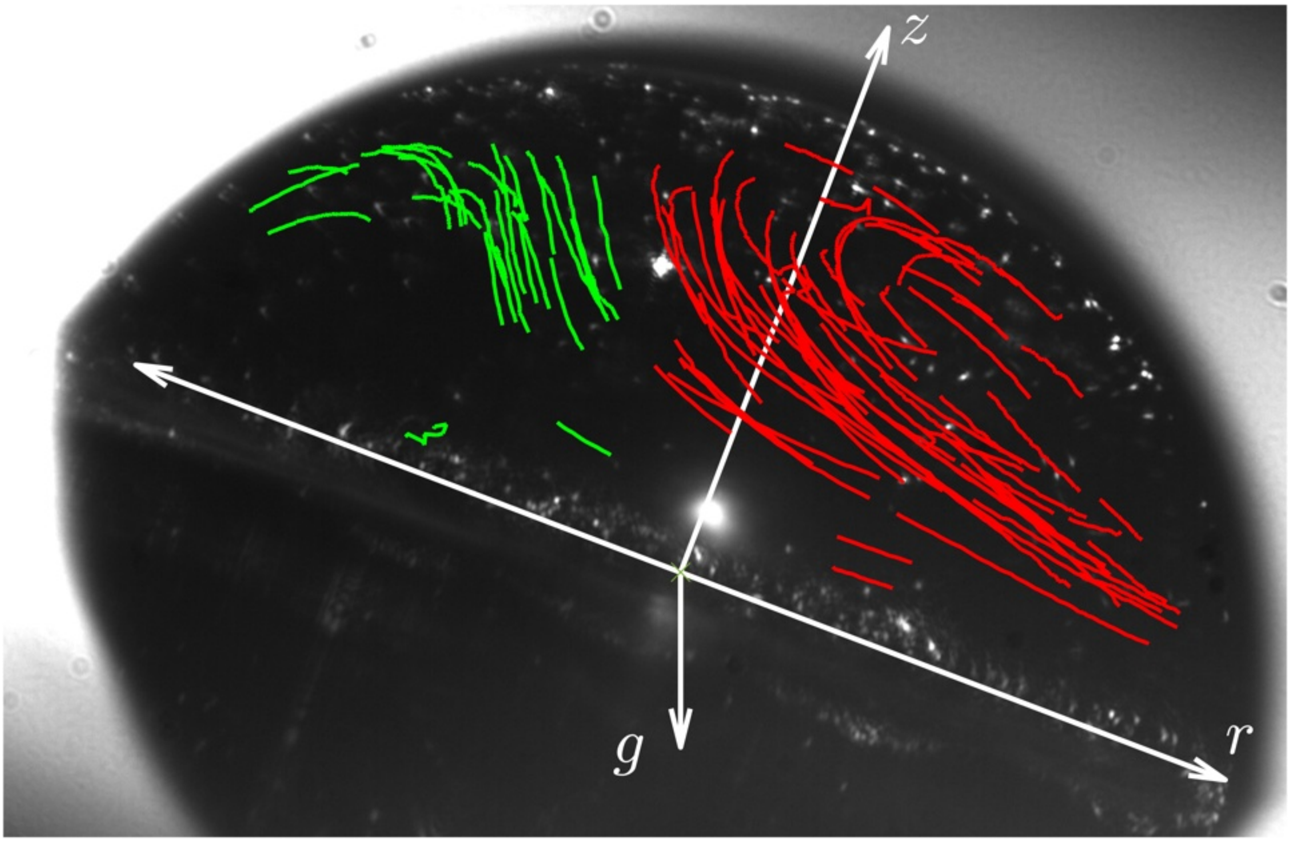


Figure S14: Particle trajectories obtained using PTV on a droplet of composition Blank A evaporated at RH = 40%. The droplet is tilted by 20° with respect to the horizontal. The green trajectories correspond to particles located to the left of the gravity vector, while the red trajectories correspond to those on the right.

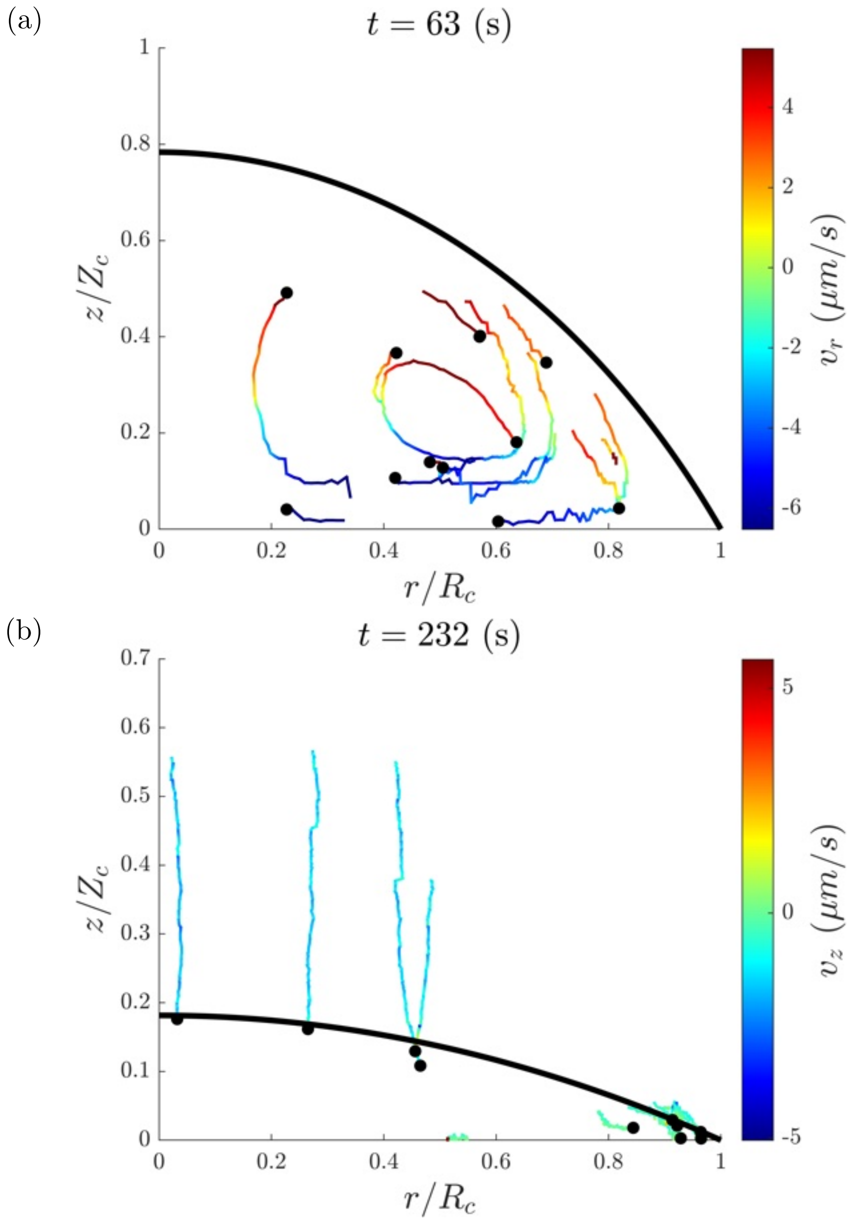


Figure S15: GDPT measurements of particle trajectories. (a) Trajectories obtained during a first stage of the droplet evaporation process. The interface has not receded enough to capture enough particles. Thus, particles remain in the bulk, where they describe the recirculatory motion studied experimentally and theoretically in the main text. (b) Particle trajectories obtained during the last stage of the droplet evaporation process. The interface has receded substantially with respect to its initial position, which has lead to the capture of the tracer particles.Drop of composition Blank A, initial volume $0.5\mu L$, evaporated at RH $= 40\%$.

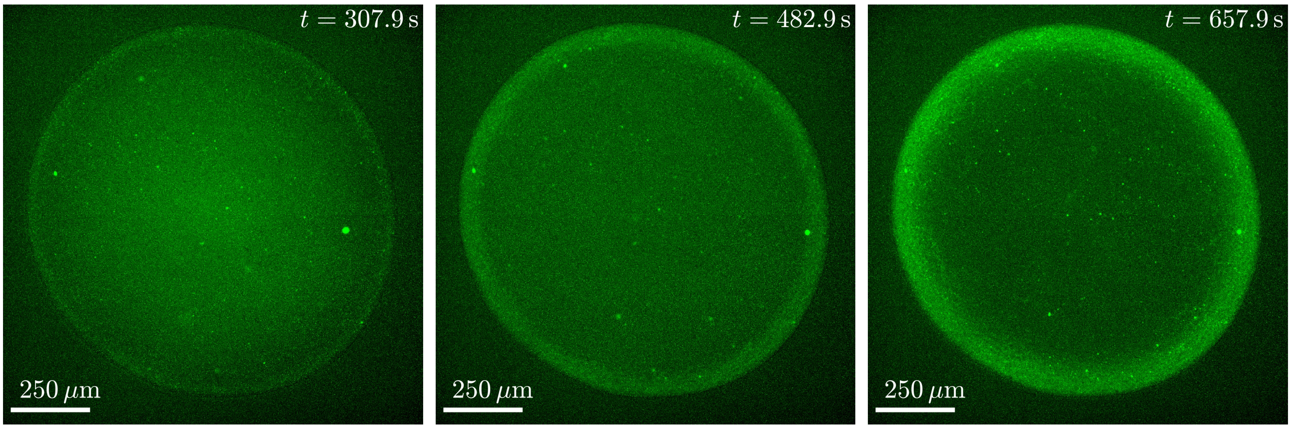


Figure S16: Subsequent fluorescent images of a droplet of composition A evaporating at RH = 70%

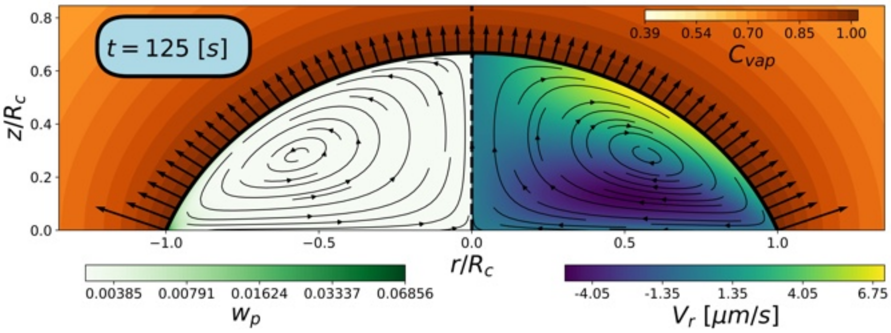


Figure S17: Simulations of a droplet of artificial saliva evaporated at RH = 40% with initial volume $V_0 = 0.81 \mu L$.

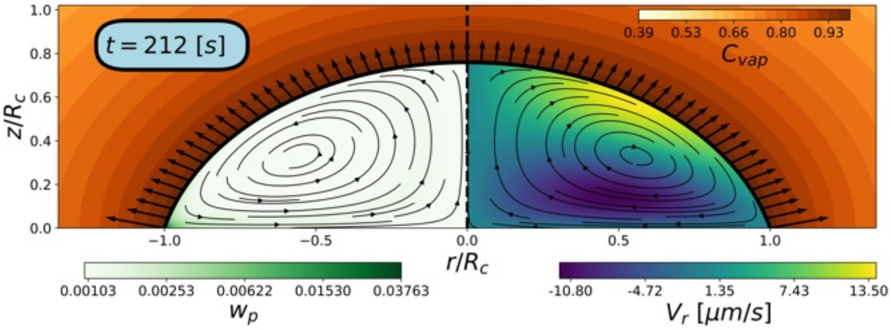


Figure S18: Simulations of a droplet of composition Blank B evaporated at RH = 40% with initial volume $V_0 = 1.5 \mu L$.

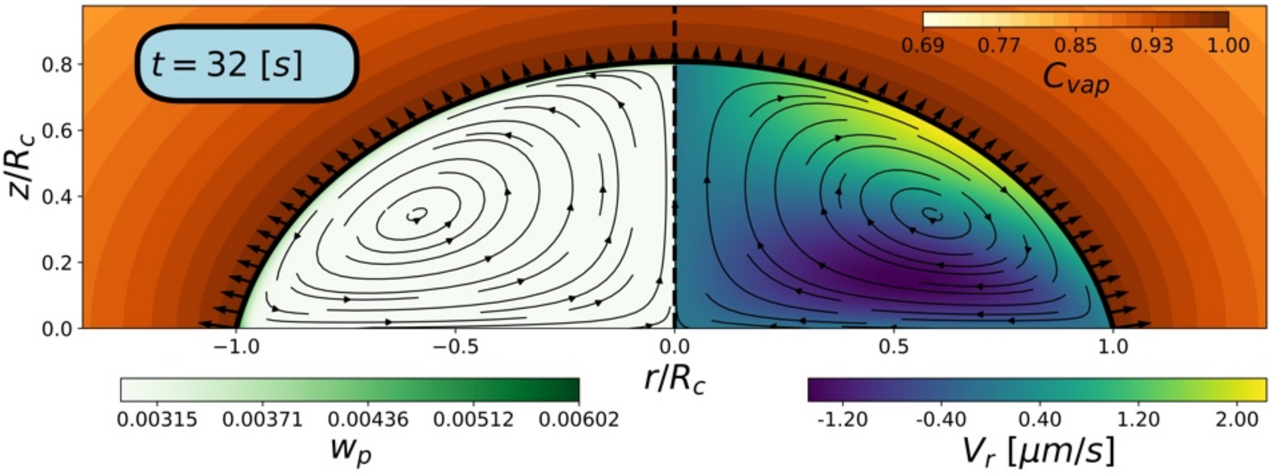


Figure S19: Simulations of a droplet of artificial saliva evaporated at RH = 70% with initial volume $V_0 = 1.8\mu L$.
